# Conformational dynamics of SARS-CoV-2 trimeric spike glycoprotein in complex with receptor ACE2 revealed by cryo-EM

**DOI:** 10.1101/2020.06.30.177097

**Authors:** Cong Xu, Yanxing Wang, Caixuan Liu, Chao Zhang, Wenyu Han, Xiaoyu Hong, Yifan Wang, Qin Hong, Shutian Wang, Qiaoyu Zhao, Yalei Wang, Yong Yang, Kaijian Chen, Wei Zheng, Liangliang Kong, Fangfang Wang, Qinyu Zuo, Zhong Huang, Yao Cong

## Abstract

The recent outbreaks of severe acute respiratory syndrome coronavirus 2 (SARS-CoV-2) and its rapid international spread pose a global health emergency. The trimeric spike (S) glycoprotein interacts with its receptor human ACE2 to mediate viral entry into host-cells. Here we present cryo-EM structures of an uncharacterized tightly closed SARS-CoV-2 S-trimer and the ACE2-bound-S-trimer at 2.7-Å and 3.8-Å-resolution, respectively. The tightly closed S-trimer with inactivated fusion peptide may represent the ground prefusion state. ACE2 binding to the up receptor-binding domain (RBD) within S-trimer triggers continuous swing-motions of ACE2-RBD, resulting in conformational dynamics of S1 subunits. Noteworthy, SARS-CoV-2 S-trimer appears much more sensitive to ACE2-receptor than SARS-CoV S-trimer in terms of receptor-triggered transformation from the closed prefusion state to the fusion-prone open state, potentially contributing to the superior infectivity of SARS-CoV-2. We defined the RBD T470-T478 loop and residue Y505 as viral determinants for specific recognition of SARS-CoV-2 RBD by ACE2, and provided structural basis of the spike D614G-mutation induced enhanced infectivity. Our findings offer a thorough picture on the mechanism of ACE2-induced conformational transitions of S-trimer from ground prefusion state towards postfusion state, thereby providing important information for development of vaccines and therapeutics aimed to block receptor binding.

## Introduction

Coronaviruses are a family of large, enveloped, positive-stranded RNA viruses that cause upper respiratory, gastrointestinal and central nervous system diseases in humans and other animals (Song et al., 2018; Walls et al., 2019). In the past few decades, new evolved coronaviruses have posed a global threat to public health, including the outbreaks of the severe acute respiratory syndrome coronavirus (SARS-CoV) in 2002-2003 and the Middle East respiratory syndrome coronavirus (MERS-CoV) in 2012 which had caused thousands of infection, and the mortality rate of them was about 10% and 34.4%, respectively (Rabaan et al., 2020). The recent coronavirus disease 2019 (COVID-19) pandemic is caused by a novel coronavirus named severe acute respiratory syndrome coronavirus 2 (SARS-CoV-2). On June 29, 2020, there had been 9,962,193 laboratory-confirmed SARS-CoV-2 infections globally, leading to 498,723 deaths. To date, there is no approved therapeutics or vaccines against SARS-CoV-2 and other human-infecting coronaviruses.

As in other coronaviruses, the spike (S) glycoprotein of SARS-CoV-2 is a membrane-fusion machine that mediates receptor recognition and viral entry into cells and is the primary target of the humoral immune response during infection (Rabaan et al., 2020; Tang et al., 2020). The S protein is a homotrimeric class I fusion protein that forms large protrusions from the virus surface and undergoes a substantial structural rearrangement to fuse the viral membrane with the host-cell membrane once binds to a host-cell receptor (Bosch et al., 2003; Li, 2016). The S protein ectodomain consists of a receptor-binding subunit S1 and a membrane-fusion subunit S2 (Tang et al., 2020; Walls et al., 2020; Wrapp et al., 2020). Two major domains in coronavirus S1 have been identified, including an N-terminal domain (NTD), and a C-terminal domain (CTD) also called receptor binding domain (RBD). Following the RBD, S1 also contains two sub-domains (SD1 and SD2) (Wrapp et al., 2020). The S2 contains a variety of motifs, starting with the fusion peptide (FP). The FP describes a short segment, conserved across the viral family and composed of mostly hydrophobic residues, which inserts in the host-cell membrane to trigger the fusion event (Epand, 2003; Tang et al., 2020). Recent cryoelectron microscopy (cryo-EM) studies on the stabilized ectodomain of SARS-CoV-2 S protein revealed a closed state of S trimer with three RBD domains in “down” conformation (Walls et al., 2020), as well as an open state with one RBD in the “up” conformation, corresponding to the receptor-accessible state (Walls et al., 2020; Wrapp et al., 2020). Unlike in MERS-CoV S protein (Pallesen et al., 2017), the two or three RBD “up” conformation has not been detected for SARS-CoV-2 S trimer.

SARS-CoV-2 S and SARS-CoV S share 76% amino acid sequence identity, yet, they bind the same host-cell receptor—human angiotensin-converting enzyme 2 (ACE2) (Hoffmann et al., 2020; Wang et al., 2020; Zhou et al., 2020). It is usually considered that the transition process towards the postfusion conformation is triggered when the S1 subunit binds to a host-cell receptor; receptor binding destabilizes the prefusion trimer, resulting in shedding of the S1 subunit and transition of the S2 subunit to a stable postfusion conformation (Walls et al., 2017b). The available crystal structures of the RBD domain of SARS-CoV-2 interacting with the extracellular peptidase domain (PD) of ACE2, together with the cryo-EM structure of RBD domain associated with the full length ACE2 provided important information on the RBD-ACE2 interaction interface, revealing that the residues S438 to Q506, known as the receptor-binding motif (RBM), within RBD directly interact with ACE2 (Lan et al., 2020; Wang et al., 2020; Yan et al., 2020). However, a complete picture of ACE2 associating with the SARS-CoV-2 trimeric S protein is still missing, and it remains elusive on how ACE2 binding induces SARS-CoV-2 S trimer conformational destabilization to facilitate transitions towards the postfusion state.

Here, we present cryo-EM structures of SARS-CoV-2 S trimer in a tightly closed state, and the S trimer in complex with the receptor ACE2 (termed SARS-CoV-2 S-ACE2) at 2.7 Å and 3.8 Å resolution, respectively, in addition to a S trimer structure in the unliganded open state. The tightly closed ground prefusion state with originally dominant population may indicate a conformational masking mechanism of immune evasion for SARS-CoV-2 spike. Our data suggested there is one RBD in the “up” conformation and is trapped with ACE2 in the S-ACE2 complex; ACE2 can greatly shift the conformational landscape of S trimer, and trigger continuous swing motions of ACE2-RBD in the context of the S trimer resulting in conformational dynamics in S1 subunits. We demonstrated the RBM T470-T478 loop and residue Y505 as viral determinants for specific recognition of SARS-CoV-2 RBD by ACE2. Our findings provide a blueprint for the understanding of the mechanisms of ACE2-induced conformational dynamics and resulted conformational transitions of the S trimer towards postfusion state, which may benefit anti-SARS-CoV-2 drug and vaccine development.

## Results

### An uncharacterized tightly closed state of SARS-CoV-2 S trimer

Prefusion stabilized ectodomain trimer of SARS-CoV-2 S glycoprotein was produced from HEK293F cells using the strategy also adopted in other studies (Fig. S1A) (Kirchdoerfer et al., 2018; Miroshnikov et al., 1998; Pallesen et al., 2017; Tortorici et al., 2019; Walls et al., 2017a; Walls et al., 2020; Walls et al., 2016; Walls et al., 2019; Wrapp et al., 2020), and was subjected to cryo-EM single-particle analysis (Fig. S2A-B). Our initial reconstruction suggested a preferred orientation problem associated with the S trimer (highly preferred “side” orientation but lacking tilted top views, Fig. S2C), which is also the case for the influenza hemagglutinin (HA) trimer (but highly preferred “top” orientation) (Tan et al., 2017). To overcome this problem, we adopted the recently developed tilt stage strategy in data collection with additional data collected at 30° and 40° tilt angles (Tan et al., 2017). This allowed us to obtain a cryo-EM structure of SARS-CoV-2 S trimer in a closed state at 2.7 Å resolution (with imposed C3 symmetry, termed S-closed) (Figs. 1A, and S2-S3, Movie 1). Excitingly, after overcoming the preferred orientation problem, our S-closed map very well resolved the peripheral edge of the NTD domain (Fig. 1A-C), which was less well resolved in the recent reports (Walls et al., 2020; Wrapp et al., 2020). This enabled us to build a more complete model of the SARS-CoV-2 S trimer containing the previously missing loop regions (including Q14-P26, K77-F79, Y144-N164, Q173-N185, R246-S247, and S255-A262, Fig.1B, S2G); additionally, the S469-C488 loop in the RBM subdomain was also captured in our structure (Fig. 1D).

**Figure 1.**
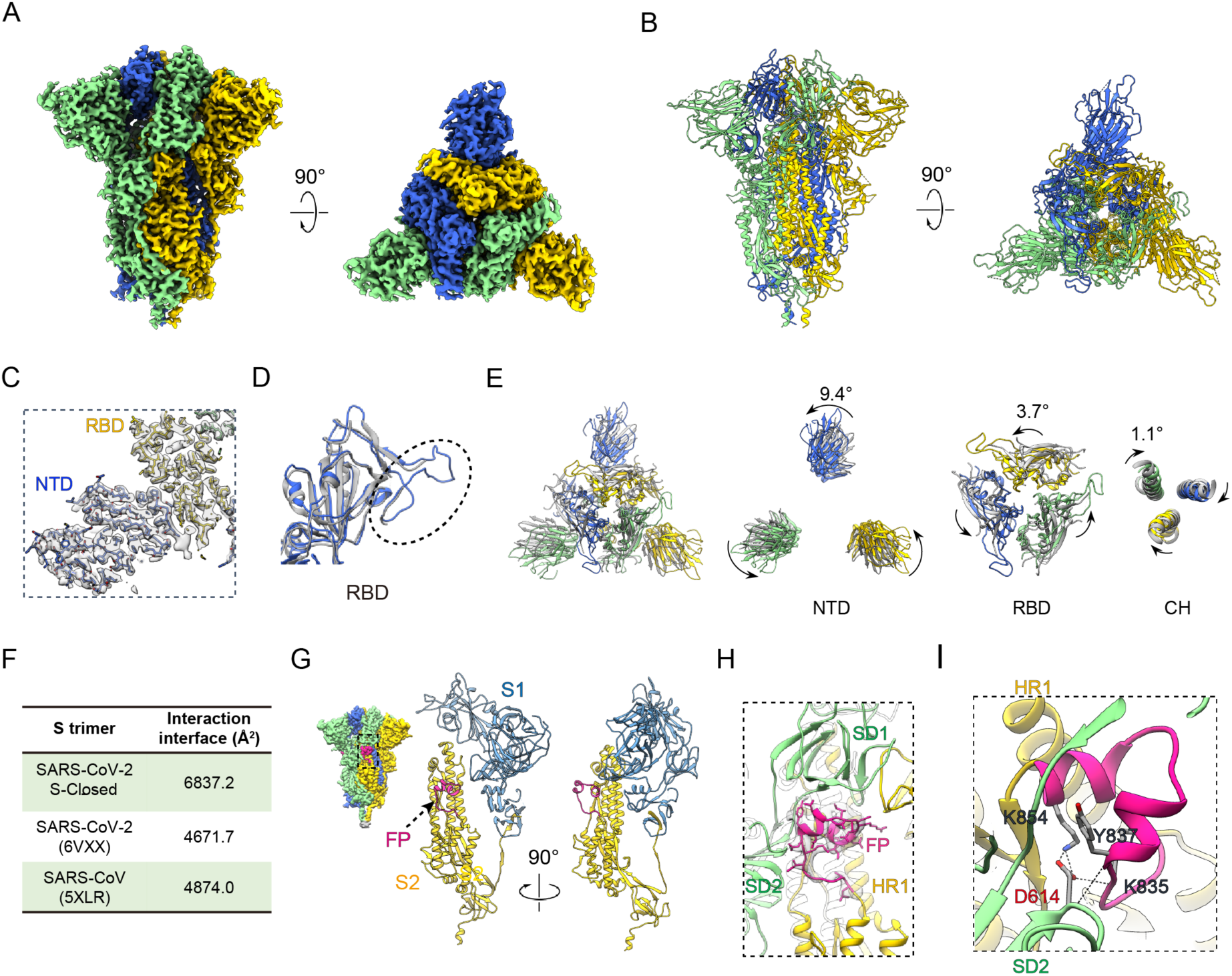
An uncharacterized tightly closed conformation of SARS-CoV-2 S trimer. (A-B) Cryo-EM map and pseudo atomic model of SARS-CoV-2 S trimer in a tightly closed state, with three protomers shown in different color. (C) Close up view of the model-map fitting in the NTD and RBD regions of S1 subunit, illustrating most of the NTD region was well resolved. (D) Overlaid structures of our S-closed structure (blue) with the recent structure of SARS-CoV-2 S in closed state (gray, 6VXX), illustrating the RBM S469-C488 loop was newly captured in our structure (highlighted in dotted ellipsoid). (E) Top view of the overlaid structures as in (D) (left panel) and zoom-in views of specific domains, showing there is a dramatic anti-clockwise rotation in S1 especially in NTD, and a slight clockwise rotation in the central CH, resulting in a twisted tightly closed conformation. (F) Protomer interaction interface analysis by PISA. (G)The location of the newly resolved FP fragment (in deep pink) within the S trimer (left) and one protomer. S1 and S2 subunit is colored steel blue and gold, respectively. (H) Model-map fitting for the newly resolved FP fragment. (I) Close up view of the interactions between D614 from SD2 and FP, with the hydrogen bonds labeled in dotted lines and the L828-F855 region in FP in deep pink.

Interestingly, compared with the recent closed state SARS-CoV-2 S trimer structure (Walls et al., 2020), our map represents an uncharacterized tightly closed conformation. For instance, the upper portion of S1 subunit especially NTD and RBD depicts an anti-clockwise rotation of 9.4° and 3.7°, respectively (Fig. 1E). Accompanying this rotation, there is a slight inward tilt leading the peripheral edge of NTD exhibiting a 12.4 Å inward movement for Cα of T124 (Fig. S2G). These motions can be propagated to the central helix (CH) of S2 subunit, generating a clockwise rotation of 1.1° (Fig. 1E). This central portion clockwise rotation associating with the outer potion opposite anti-clockwise rotation in reality twists the complex in a more compact conformation. Indeed, the average interaction interface between protomers increased from ∼4,671.7 Å^2^ in their structure to 6,837.2 Å^2^ in our structure (Fig. 1F). Taken together, our map represents a tightly closed state of the SARS-CoV-2 S trimer, not captured before. Furthermore, when comparing our SARS-CoV-2 S-closed structure with the closed state SARS-CoV S trimer cryo-EM structure (Gui et al., 2017), there is an anti-clockwise rotation of 10.5° and 8.7° in NTD and RBD, respectively, and a clockwise rotation of 4.3° in CH region from their structure to our S-closed structure, associating with a RBD inward shift towards the central axis (RMSD of 6.7 Å, Fig. S2H). Collectively, our S-closed structure appears more compact than that of SARS-CoV S trimer (6,837.2 Å^2^ vs. 4,874.0 Å^2^ in interaction interface, Fig. 1F). Altogether, our study revealed a tightly closed conformation of SARS-CoV-2 S trimer, not observed in the homologous SARS-CoV S neither, extending the detected conformational space of SARS-CoV-2 spike protein.

### The tightly closed state with stably packed fusion peptide may represent the ground prefusion state of SARS-CoV-2 S trimer

The hydrophobic fusion peptide, immediately after the S2’ cleavage site and essential for host-cell membrane fusion, is highly conserved among SARS-CoV-2, SARS-CoV, and MERS-CoV S proteins (Tang et al., 2020). Still, the majority of FP is missing in the available SARS-CoV-2 S trimer structures. Thus, how it folds and where it locates within S trimer of the virus and how it can be activated remain unclear. Here, our S-closed map enabled us to capture the entire FP of SARS-CoV-2 including the previously undetected L828-Q853 fragment, which locates on the flank surface of S trimer, surrounded by HR1 of S2 subunit from the same protomer, and SD1/SD2 of S1 subunit from the clockwise neighboring protomer (Fig. 1G-H). The FP fragment is well ordered, forming two small helixes (Y837-G842, L849-F855) and connecting loops (Fig. 1G-H). This observation further substantiates the notion that our S-closed structure with inactivated FP most likely represents the ground prefusion state.

Further interaction analysis revealed that SD2 and HR1 can form hydrogen bonds/salt bridges with the FP fragment, and SD2 plays a key role in this interaction involving in 6 predicted hydrogen bonds/salt bridges (Table S2). Noteworthy, among the 6 SD2-FP interactions, D614 from SD2 contributes to the formation of 4 hydrogen bonds/salt bridges, majorly through its sidechain atoms, with K835, Y837 and K854 of FP, suggesting D614 may be essential in the interaction with and stabilization of FP (Fig. 1I and Table S2). This could be related to the recent reports suggesting that the D614G mutation of SARS-CoV-2 S enhanced viral infectivity (more in discussion) (Korber et al., 2020). Interestingly, it appears that before being activated, FP could serve as a linkage that wraps around the neighboring protomers in their S1/S2 interface and simultaneously connects S1 with S2, this way to coordinately lock the S trimer in the tightly closed ground prefusion state (Fig. 1G-H).

Moreover, in this dataset the dominant population of the particles (∼94%) is in the tightly closed state; although performed multiple rounds of 3D classification, eventually we found only a minor population (6%) of the particles is in the open state (Fig. S3). Our observations indicate that the open state SARS-CoV-2 S might be intrinsically dynamic and only exist transiently to expose the RBD domain. Interestingly, the dominant population of the SARS-CoV-2 S trimer is in the ground prefusion state with inactivated FP and all the RBD domains buried, which may result in “conformational masking” preventing antibody binding and neutralization, similar to that described for HIV-1 envelope (Env) (Kwong et al., 2002; Munro et al., 2014). The population distribution of closed and open state of SARS-CoV-2 S varies among different studies (Walls et al., 2020; Wrapp et al., 2020), which is reminiscent of observations made with SARS-CoV S and MERS-CoV S trimers. This observed variation could be potentially due to subtle difference in chemical condition used by different research groups (Gui et al., 2017; Kirchdoerfer et al., 2018; Pallesen et al., 2017; Song et al., 2018; Walls et al., 2019; Yuan et al., 2017).

### A complete architecture of the SARS-CoV-2 S trimer in complex with ACE2

To gain a thorough picture on how the receptor ACE2 binding induces conformational dynamics of the SARS-CoV-2 S trimer and triggers transition towards the postfusion state, we determine the cryo-EM structure of SARS-CoV-2 S trimer in complex with human ACE2 PD domain to 3.8 Å resolution (termed SARS-CoV-2 S-ACE2, Figs. 2A, S4A-E, and S5). Further focused-refinements improved the resolution of the S trimer portion of the map to 3.3 Å, and the connectivity in the ACE2-RBD portion of the map, respectively (Fig. S4E, S5). We then built a pseudo atomic model of the complex with combined map information (Fig. 2B). To the best of our knowledge, the structure of SARS-CoV-2 S-ACE2 complex has not been reported before. In this dataset we additionally captured an unliganded S trimer in the open state with one RBD up (resolved to 6.0 Å resolution, termed S-open), but did not detect the closed state (Fig. S4D-F and S5). We should mention that our bio-layer interferometry (BLI) assay revealed a relatively rapid disassociation kinetics between ACE2 and the S trimer (*k*_*off*_ = 4.56×10^−3^ s^-1^, Fig. S1E). We thus determined the complex structure in the presence of trace amount of cross linker glutaraldehyde (Methods). Additionally, we also determined the S-ACE2 complex structure without cross linker at 5.3 Å resolution, and the two maps are in comparable conformation, suggesting that addition of cross linker did not change the conformation of the complex (Fig. S4G). We then used the S-ACE2 map at 3.8 Å resolution for detailed structural analysis.

**Figure 2.**
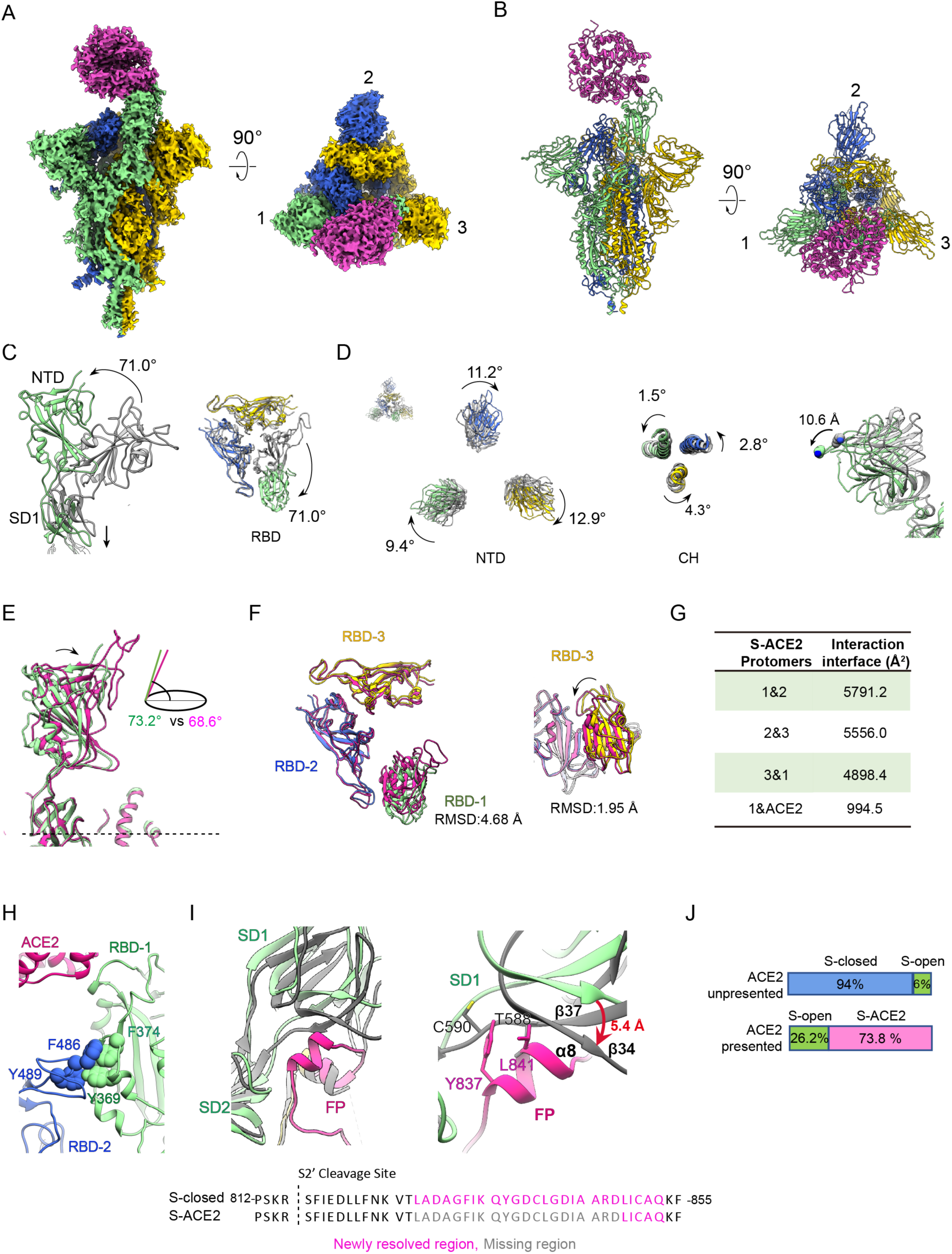
A complete architecture of the SARS-CoV-2 S-ACE2 complex. (A-B) Cryo-EM map and pseudo atomic model of SARS-CoV-2 S-ACE2 complex. We named the RBD up protomer as protomer 1 (light green), and the other two RBD down ones as protomer 2 (royal blue) and protomer 3 (gold). ACE2 was colored in violet red. (C) Side and top views of the overlaid S-open (color) and S-closed (dark grey) structures, showing in the open process there is a 71.0° upwards/outwards rotation of RBD associated with a downwards shift of SD1 in protomer 1. (D) Rotations of NTD and CH from S-closed (grey) to S-open (in color) state, with the NTD also showing a downwards/outwards movement. (E) Side view of the overlaid S-ACE2 (violet red) and S-open (light green) protomer 1 structures, showing the angle between the long axis of RBD and the horizontal plane of S trimer reduces from the S-open to the S-ACE2 state. (F) Top and side views of the overlaid S-ACE2 (violet red) and S-open (color) RBD structures, showing the coordinated movements of RBDs. (G) Protomer interaction interface analysis of S-ACE2 by PISA. (H) Aromatic interactions between the core region of the up RBD-1 (green) and the RBM T470-F490 loop of the neighboring RBD-2 (blue). (I) Overlaid structures of S-ACE2 (grey) and S-closed (color, with the FP fragment in deep pink), indicating a downwards shift of SD1 and most of the FP is missing in S-ACE2 state. Close up view (right panel) of the potential clashes between the downwards shifted SD1 β34 and α8 helix of FP. (J) Population shift between the ACE2-unpresented and ACE2-presented S trimer samples.

To inspect the conformational changes from the closed state to the unliganded open state, we first overlaid our S-open with our S-closed structures together. In the S-open structure, the only up RBD domain from protomer 1 (termed RBD-1) shows a 71.0° upwards/outwards rotation, resulting in an exposed RBM region accessible for ACE2 binding (Fig. 2C). This RBD-1 rotation can be propagated to the underneath SD1, inducing a downwards movement of SD1 (Fig. 2C). We also noticed a considerable clockwise rotation of 9.4°, 11.2°, and 12.9° in NTD for protomer 1, 2, and 3, respectively, and anti-clockwise rotations in the CH of corresponding S2 subunit, greatly untwisting the S trimer from the tightly closed state (Fig. 2D). Associated with this S1 untwisting, there is a downwards/outwards movement of NTDs in the scale of ∼10 Å (Fig. 1D, right panel). These combined untwisting motion could release the original protomer interaction strength, beneficial for the transient raising up of the RBD. Moreover, our local resolution analysis on the S-open map also suggested that other than RBD-1, the consecutive RBD-2 also exhibits considerable dynamics (Fig. S4D).

Our SARS-CoV-2 S-ACE2 structure revealed that the S trimer binds with one ACE2 through the only up RBD domain, while the other two RBDs remain in the down conformation (Fig. 2A-B), suggesting ACE2 binding to SARS-CoV-2 strictly requires the up conformation of RBD. Unlike the observations made with SARS-CoV and MERS-CoV S trimers, we did not detect S trimer with two RBD domains up with bound ACE2 (Kirchdoerfer et al., 2018; Pallesen et al., 2017). Though our S-ACE2 and S-open structures generally resemble each other especially in the S2 region, there are noticeable differences in the S1 region. Specifically, after ACE2 binding, the up RBD-1 from the S-open state can be pushed tilting downwards slightly, with the angle to the horizontal plane of S trimer reduced from 73.2° to 68.6° in ACE2 bound state (Fig. 2E). This ACE2 binding induced motion of RBD-1 could be propagated to the neighboring RBD-2 and the consecutive RBD-3 (RMSD: 1.95 Å, Fig. 2F), collectively disturbing the allosteric network of the fusion machinery. Indeed, the neighboring protomer interaction interface was reduced from the original ∼6837.2 Å^2^ in the S-closed state to 4898.4∼5791.2 Å^2^ in the ACE2 bound state (Fig. 2G). Altogether, these S1 subunits untwisting and RBD-1 tilting motions could destabilize the prefusion state of S trimer, prepared for the subsequent conformational transitions towards the postfusion state.

Interestingly, our S-ACE2 structure showed that the core region of the up RBD-1 and the RBM T470-F490 loop of the neighboring RBD-2 could form aromatic interactions with the involvement of Y369/F374 from RBD-1 and F486/Y489 from RBD-2 (Fig. 2H), potentially enhancing interactions between neighboring S1 subunits, thus beneficial for subsequent simultaneous release of S1 subunits. This interaction was not detected in the counterpart of the homologous SARS-CoV S-ACE2 structure, likely due to longer distance between the adjacent “up” and “down” RBDs in that structure (Kirchdoerfer et al., 2018; Song et al., 2018).

Noteworthy, the originally stably packed FP from protomer 3 surrounded by SD1/SD2 of the neighboring protomer 1 in the S-closed structure is now mostly missing in the S-ACE2 structure, which is also the case in the S-open structure. This is mostly caused by the S trimer untwisting-motion induced downwards shift of SD1 (Fig. 2C, I). Indeed, the β34/β37 strands within SD1 shift downwards for up to 5.4 Å; consequently, the C590 and T588 from β37 and the connecting loop could clash with the Y837 and L841 of the originally packed α8 helix of FP (Fig. 2I), potentially resulting in destabilization and activation of the FP motif from protomer 3. Since the untwisting/downwards-shift motions of S1 subunits are allosterically coordinated within the S trimer in its opening process, the density corresponding to FPs in protomer 1 and 2 are also missing, indicating a coordinated activation mechanism of FP, which may be one of the key elements prepared for the subsequent fusion of S trimer.

Strikingly, our data further suggested that the presence of ACE2 could greatly shift the population landscape of S trimer, i.e. from the original 94% of closed prefusion state and 6 % fusion-prone open state in the absence of ACE2, to 26.2% unliganded open state and 73.8% ACE2 bound open state in the ACE2 present sample (Fig. 2J). Therefore, in the presence of ACE2, the open state S trimer (despite this state only exits transiently with minor population in the absence of ACE2) interacts with ACE2, and such interaction could break the balance between particle populations and greatly shift the S trimer conformational landscape towards the open state, favorable for the receptor binding and transitions towards postfusion state.

### The T470-T478 loop and residue Y505 within RBM play vital roles in the engagement of SARS-CoV-2 spike with host-cell receptor ACE2

According to our SARS-CoV-2 S-ACE2 cryo-EM structure, the overall ACE2-RBD interaction interface is comparable to that of the crystal structures of the RBD domain of SARS-CoV-2 S interacting with the ACE2 PD domain (Fig. 3A) (Lan et al., 2020; Wang et al., 2020), i.e. our structure revealed 19 residues of RBD are in contact with 17 residues of ACE2 with a distance cut-off of 4 Å (Table S3). Sequence alignment demonstrated that the RBM T470-F490 loop is the most diversified region between SARS-CoV-2 and SARS-CoV S proteins (Fig. S6). In line with this, structural comparison revealed that the conformation of the RBM T470-F490 loop in our SARS-CoV-2 S-ACE2 structure is very distinct from that in the SARS-CoV RBD-ACE2 crystal structure (Fig. 3B) (Li et al., 2005). Noteworthy, the RBM T470-F490 loop can originally be resolved in our S-closed structure, but is mostly missing in our S-open structure, indicating the T470-F490 loop may be activated in the open state. In our S-ACE2 structure, a portion of this loop forms contact with the N-terminal helix of ACE2 (Fig. 3A), for instance, A475 within this loop could interact with S19/T27 of ACE2 (Table S3), suggesting that the RBM T470-F490 loop may play an important role in receptor recognition. Moreover, the S-ACE2 structure indicated that the Q498-Y505 region located in the other edge of RBM could also form close contact with ACE2, i.e. Y505 could form hydrogen bonds/contacts with A386/R393/K353/G354 from ACE2.

**Figure 3.**
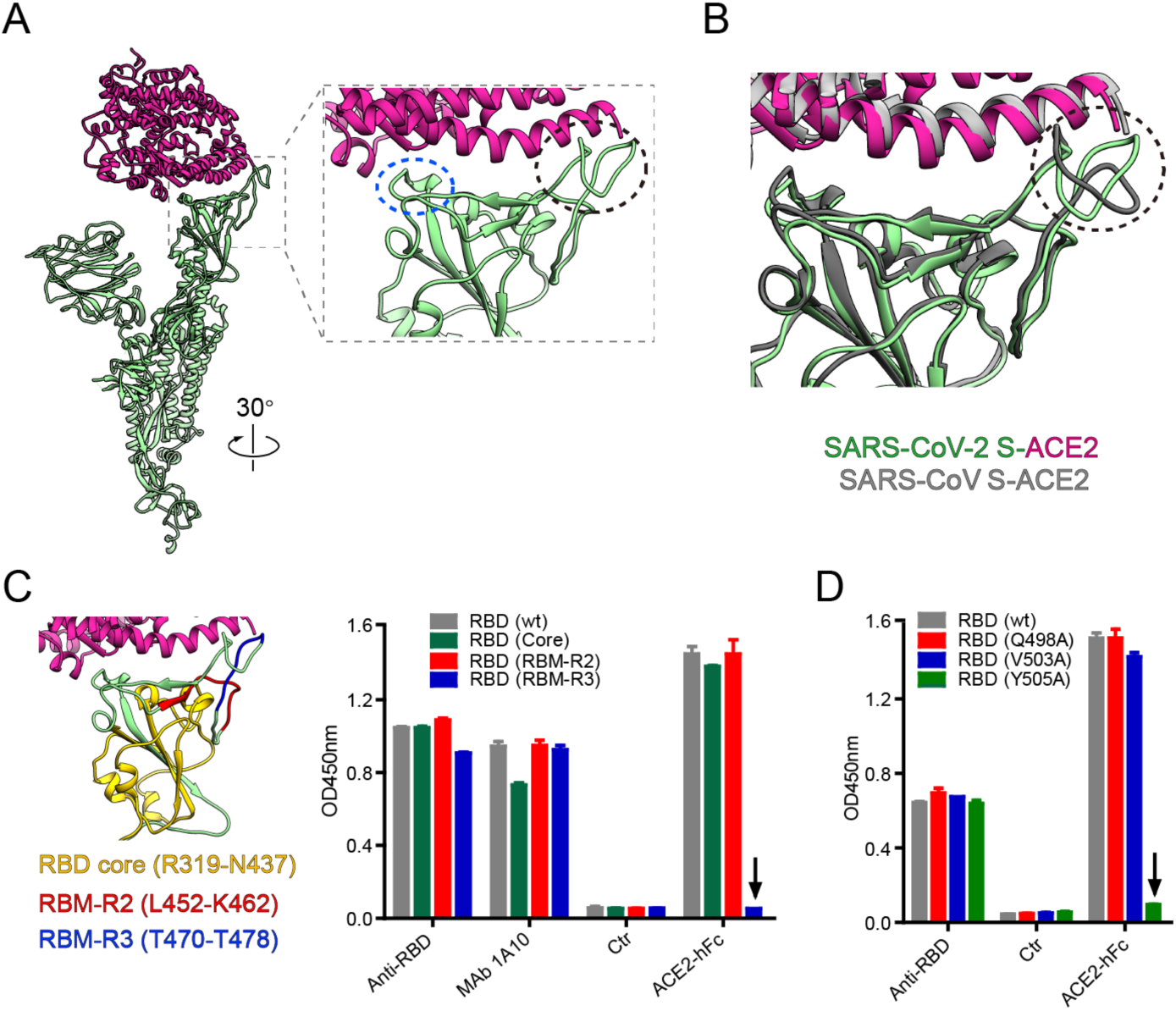
The T470-T478 loop and residue Y505 within RBM play vital roles in the engagement of SARS-CoV-2 spike with receptor ACE2. (A) The overall view of ACE2 (violet red) bound protomer 1 (light green) from our S-ACE2 structure, and zoom in view of the interaction interface between ACE2 and RBD, with the key contacting elements T470-F490 loop and Q498-Y505 within RBM highlighted in black ellipsoid and blue ellipsoid, respectively. (B) Superposition of our SARS-CoV-2 S-ACE2 structure with the crystal structure of SARS-CoV RBD-ACE2 (PDB: 2AJF), suggesting the RBM T470-F490 loop has obvious conformational variations. (C) Binding activities of ACE2-hFc fusion protein to wild-type (wt) and mutant SARS-CoV-2 RBD proteins determined by ELISA. Different structural elements of RBD were colored in the left panel. Anti-RBD sera and a cross-reactive MAb 1A10 served as positive controls. Ctr, an irrelevant antibody. The black arrow indicates that mutations in the RBD (RBM-R3) mutant significantly reduced the binding of ACE2-hFc compared to wild-type RBD. (D) Binding of ACE2-hFc fusion protein to wt and single-point mutant forms of SARS-CoV-2 RBD protein measured by ELISA. RBD (Q498A), RBD (V503A), and RBD (Y505A), RBD residues Q498, V503, and Y505 were mutated to Ala, respectively. The downward arrow indicates that the mutation at Y505 completely abolished the binding of ACE2 to RBD protein.

To further define the subdomains/residues critical for RBD binding to ACE2, we designed and produced three SARS-CoV-2 RBD mutant proteins, each of which had a single subdomain substituted with the counterpart of SARS-CoV. These RBD mutants were termed RBD-(Core), RBD-(RBM-R2) and RBD-(RBM-R3), which harbored R319 to N437 of the core region, L452 to K462, and T470 to T478 of the RBM from SARS-CoV, respectively (Figs. 3C and S6). Results from ACE2-binding enzyme linked immunosorbent assay (ELISA) showed that the binding activity of the three RBD mutants towards anti-RBD polyclonal antisera and the cross-reactive monoclonal antibody 1A10 was comparable to that of the wildtype SARS-CoV-2 RBD protein (Fig. 3C), indicating that the mutations did not significantly affect the overall conformation of the RBDs. The mutants RBD-(Core) and RBD-(RBM-R2) bound ACE2 as efficiently as the wildtype RBD; in contrast, RBD-(RBM-R3) completely lost ACE2-binding (Fig. 3C). These results pinpoint the RBM-R3 region (residues 470-TEIYQAGST-478) as the critical viral determinant for specific recognition of SARS-CoV-2 RBD by the ACE2 receptor. Additionally, we constructed three single-point mutants of SARS-CoV-2 RBD protein, RBD (Q498A), RBD (V503A), and RBD (Y505A). Our ELISA ACE2-binding assay showed that the mutation Y505A was sufficient to completely abolish the binding of ACE2, while the other two mutations did not show such effect (Fig. 3D), demonstrating that the residue Y505 of SARS-CoV-2 RBD is a key amino acid required for ACE2 receptor binding.

### ACE2 binding induces continuous swing motions of ACE2-RBD in the context of SARS-CoV-2 S trimer

Our SARS-CoV-2 S-ACE2 map showed well defined density for the S trimer region, but relatively lower local resolution in the associated ACE2-RBD region (Fig. S4C), suggesting considerable conformational heterogeneity of ACE2-RBD as well as relative dynamics between ACE2-RBD and the remaining part of S trimer with respect to each other. This is in line with the report showing that in SARS-CoV S trimer, the associated ACE2-RBD is relatively dynamic, showing three major conformational states with the angle of ACE2-RBD to the surface of S trimer at ∼51°, 73°, and 111°, respectively (Song et al., 2018). To better delineate the conformational space of the ACE2 engaged SARS-CoV-2 S trimer, we performed multi-body refinement in Relion 3.1 (Fernandez-Leiro and Scheres, 2017).

Principal component analysis of the movement revealed that approximately 68% of the movement of the complex is described by the first three eigenvectors representing swing motions in distinct directions relative to the S trimer (Fig. 4A). Eigenvector 1 describes a swing motion of ACE2-RBD towards RBD-2 direction with the angular range of 12.2°, eigenvector 2 corresponds to the swing motion of ACE2-RBD towards the original location of RBD-1 with the angular range of 11.9°, and eigenvector 3 describes the swing motion of ACE2-RBD along the NTD-1 to NTD-3 direction with the angular range of 9.8° (Fig. 4B). Histograms of the amplitudes along the three eigenvectors are unimodal, indicative of continuous motions (Fig. 4C). As the dynamic motions in the complex are formed by linear combination of all eigenvectors, these data suggested that ACE2-RBD processes on top of the S trimer in a non-correlated manner. Moreover, multi-body analysis on the non-cross-linked SARS-CoV-2 S-ACE2 data showed similar swing motions (Fig. S4H), indicating the presence of cross linker did not disturb the mode of ACE2-RBD motions within the S trimer. Additionally, compared with the homologous SARS-CoV S-ACE2 complex, which shows discrete movements of ACE2-RBD in one direction (similar to our eigenvector 2 direction) (Song et al., 2018), ACE2 binding to SARS-CoV-2 S induces more complex combined continuous swing motions of ACE2-RBD within the complex.

**Figure 4.**
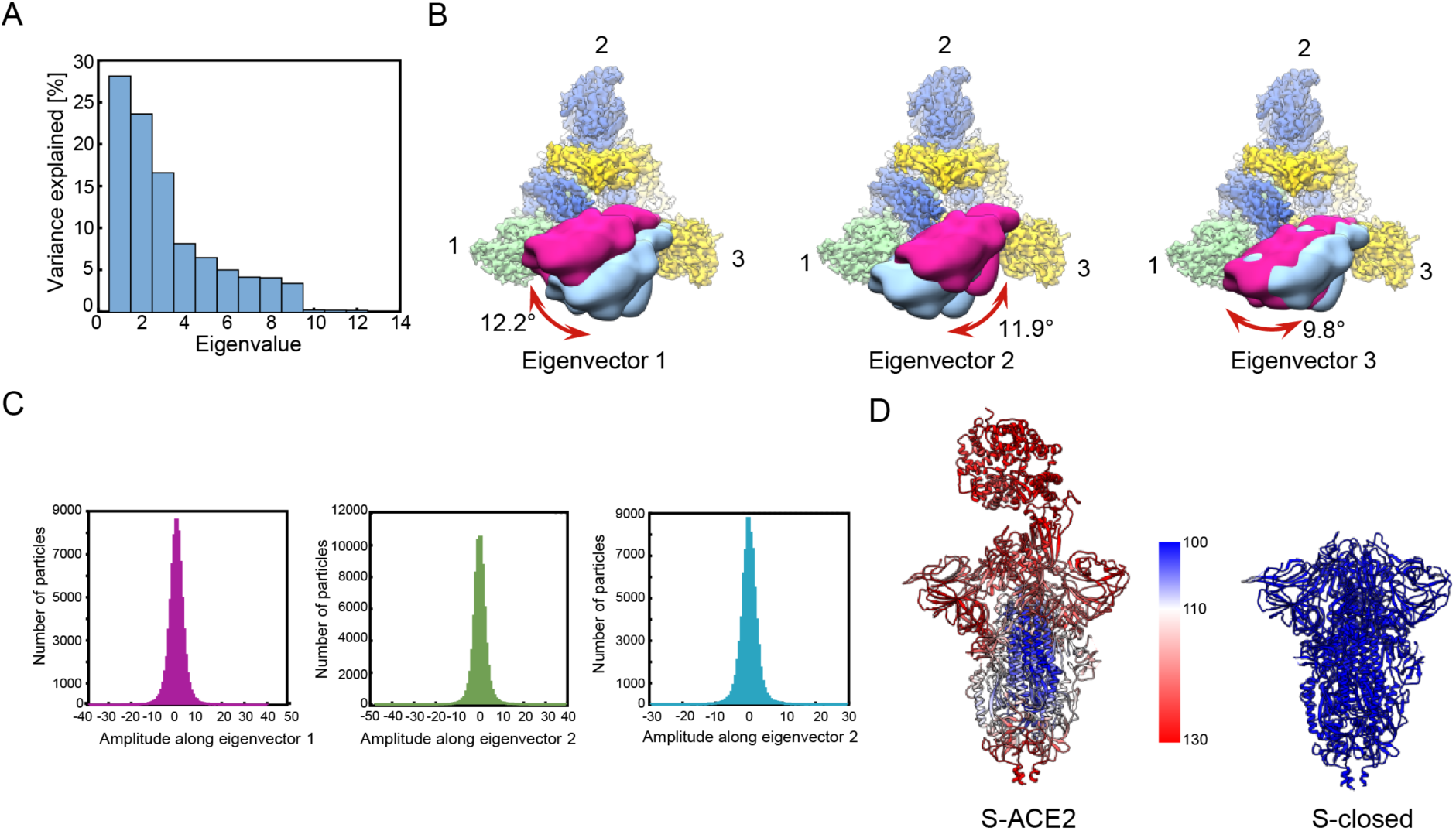
ACE2 binding induced conformational dynamics of the SARS-CoV-2 S-ACE2 complex determined by multi-body refinement. (A) The contributions of all eigenvectors to the motion in the S-ACE2 complex, with eigenvectors 1-3 dominant the contributions. (B) Top view of the map showing the three swing motions of the first 3 eigenvectors, with S trimer following the color schema as in Fig. 2, and the two extreme locations of ACE2 illustrated in deep pink and light blue densities. The swing angular range and direction are indicated in dark red arrow. (C) Histograms of the amplitudes along the first 3 eigenvectors. (D) Atomic models of S-ACE2 and S-closed, colored according to the B factor distribution (ranging from 100Å^2^ [blue] to 130Å^2^ [red]).

Putting together, our observations suggest that ACE2 receptor binding to SARS-CoV-2 S triggers considerable conformational dynamics in S1 subunits that could destabilize the prefusion S trimer. Indeed, the B-factor distribution of our S-ACE2 complex demonstrated that ACE2 binding induces strikingly enhanced dynamics in the S1 region including RBD and NTD domains (Fig. 4D), facilitating the release of the associated ACE2-S1 component and transitions of the S2 subunit towards a stable postfusion conformation. Indeed, we found a notable drop in the interaction surface between S1 and S2 subunits from the S-closed state (8982.3 Å^2^) to the S-ACE2 state (6521.7 Å^2^).

### The SARS-CoV-2 S glycan shields

It has been suggested that the large number of N-linked glycans covering the surface of the spike protein of SARS-CoV and MERS-CoV could pose challenge to antigen recognition, thus may help the virus evade immune surveillance (Walls et al., 2019; Yuan et al., 2017). Similar to SARS-CoV S, SARS-CoV-2 S also comprises 22 N-linked glycosylation, with 13 glycans in the S1 subunit and the other 9 in the S2 subunit (Fig. 5A) (Walls et al., 2020; Watanabe et al., 2020). In our S-closed structure, we resolved the density for 18 N-linked glycans per protomer (Fig. 5A-B and S2I), including two undetected glycans at site N17 and N149 located in the NTD (Fig. 5B), while the three glycans located in the flexible C-terminal region are missing as in other studies (Walls et al., 2020; Wrapp et al., 2020). Similar to MERS-CoV and SARS-CoV S trimers (Walls et al., 2017b; Walls et al., 2019; Yang et al., 2015), SARS-CoV-2 S trimer also forms a glycan hole at proximity of the S1/S2 cleavage site and the fusion peptide (near the S2’ cleavage site, Fig. 5B). Although there is an extra glycan at N657 site near the S1/S2 cleavage site in SARS-CoV-2 S, the hole region is still more sparsely glycosylated than the rest of the protomer. This glycan hole might be important for permitting the access of activating host proteases and for allowing membrane fusion to take place without obstruction (Walls et al., 2017b; Walls et al., 2019; Yang et al., 2015). Moreover, after ACE2 binding, our S-ACE2 structure revealed that the density corresponding to glycan at N165 site is weaker in protomer 1, while the other resolved glycans in the S-closed state can also be visualized in the S-ACE2 structure (Fig. 5C).

**Figure 5.**
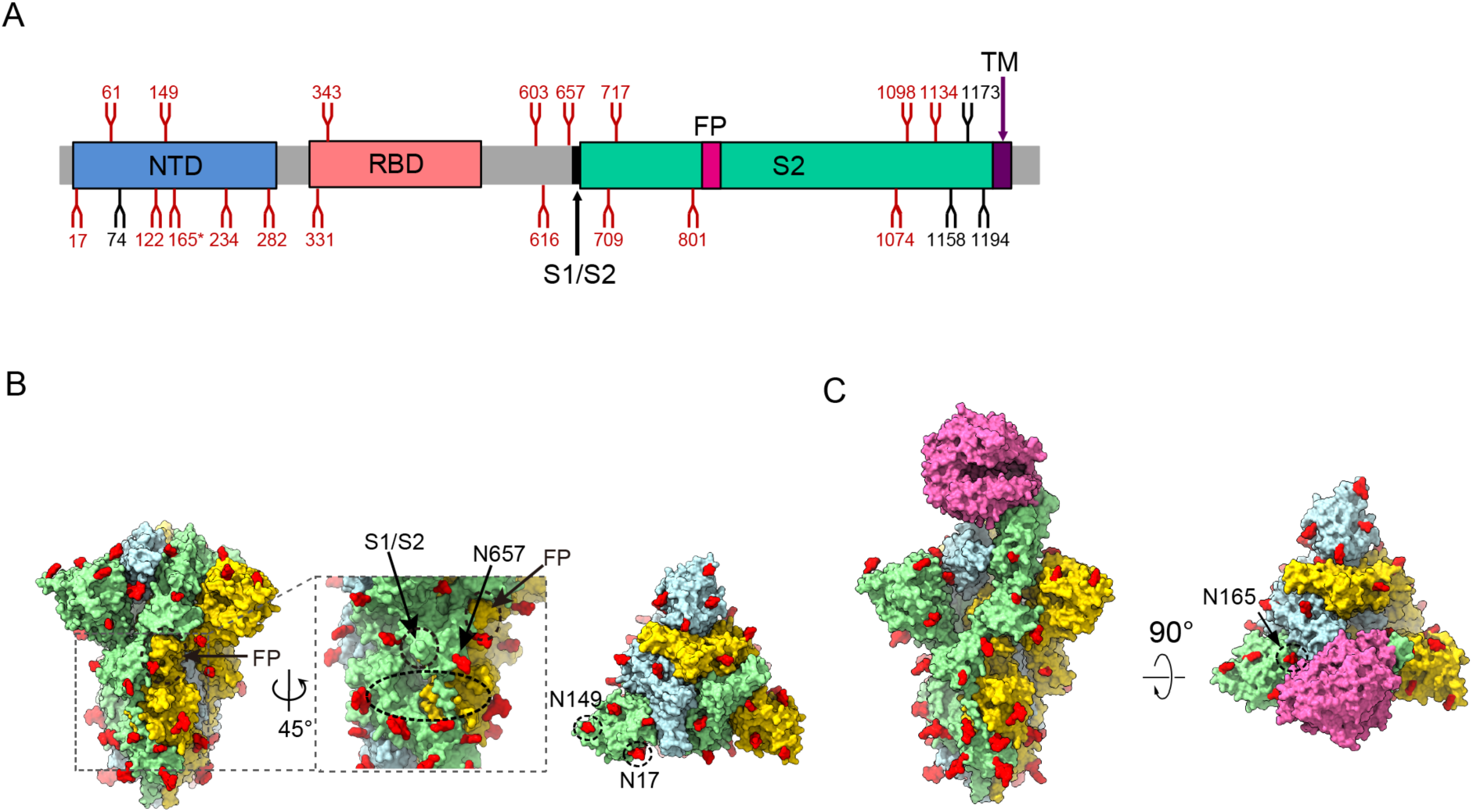
Organization of the resolved N-Linked glycans of SARS-CoV-2 S trimer. (A) Schematic representation of SARS-CoV-2 S glycoprotein. The positions of N-linked glycosylation sequons are shown as branches. 18 N-linked glycans detected in our cryo-EM map of the S-closed state are shown in red, the remaining undetected ones in black. After ACE2 binding, the glycan density that appears weaker is indicated (*). (B) Surface representation of the glycosylated SARS-CoV-2 S trimer in the S-closed state with N-linked glycans shown in red. The location of glycan hole is indicated in black doted ellipsoid, with the locations of S1/S2 and FP, and glycan at N657 site near the glycan hole indicated. The newly captured glycans at N17 and N149 sites are indicated in the top view. (C) Surface representation of the glycosylated S-ACE2 complex with N-linked glycans in red. After ACE2 binding, the glycan density that appears weaker is indicated.

## Discussion

The outbreak of COVID-19 caused by SARS-CoV-2 virus has become pandemic. Several structures of SARS-CoV-2 spike RBD domain bound to ACE2 have been reported (Lan et al., 2020; Wang et al., 2020; Yan et al., 2020). However, the complete architecture of SARS-CoV-2 trimeric S in complex with ACE2 remains unavailable, leading to an incomplete understanding of the nature of this interaction and of the resulted conformational transitions of the S trimer towards postfusion and virus entry. In the present study, we determined an uncharacterized tightly closed state of SARS-CoV-2 S trimer revealing the stably packed fusion peptide, most likely representing a previously undetected ground prefusion state of S trimer. The tightly closed S trimer with originally dominant population may indicate a conformational masking mechanism of immune evasion for SARS-CoV-2 spike. Importantly, we captured the complete architecture of SARS-CoV-2 S trimer in complex with ACE2. We found the presence of ACE2 could dramatically shift the conformational landscape of the S trimer, and after engagement the continuous swing motions of ACE2-RBD in the context of the S trimer could generate considerable conformational dynamics in S1 subunits resulting in a significant decrease in S1/S2 interface area. Furthermore, our structural data combined with biochemical analysis revealed that the RBM T470-T478 loop and residue Y505 play vital roles in the binding of SARS-CoV-2 RBD to ACE2 receptor. Our findings depict a new role of FP in stabilizing S trimer and the mechanism of FP activation, expand the detected conformational space of the S trimer, and provide structural basis on the SARS-CoV-2 spike D614G mutation induced enhanced infectivity.

Based on the data, we put forward a mechanism of ACE2 binding-induced conformational transitions of SARS-CoV-2 S trimer from the tightly closed ground prefusion state transforming towards the postfusion state (Fig. 6). In the receptor-free SARS-CoV-2 S, the majority of the S trimers is in the tightly closed ground prefusion state with inactivated FP, and only a minor population of the particles is in the transient open state with one RBD up representing the fusion-prone state, forming a dynamic balance between the two states (step 1). However, the presence of ACE2 and subsequent trapping of the RBD (discussed later) could overcome the energy barrier, break the balance and shift the conformational landscape towards the open state with an untwisting/downwards-shift motion of the S1 subunits, leading to unpacked/activated FPs, weakened interactions among the protomers, and an up RBD. In step 2, once the receptor ACE2 grasp the up RBD, the RBD will be trapped in the up conformation, and the associated ACE2-RBD together shows combined continuous swing motions on the topmost surface of the S trimer. These motions and dynamics could disturb the allosteric network and release the constrains imposed on the fusion machinery, beneficial for the releasing of the ACE2-S1 component, thereby allowing the S trimers to refold and fuse the viral and host membranes (step 3).

**Figure 6.**
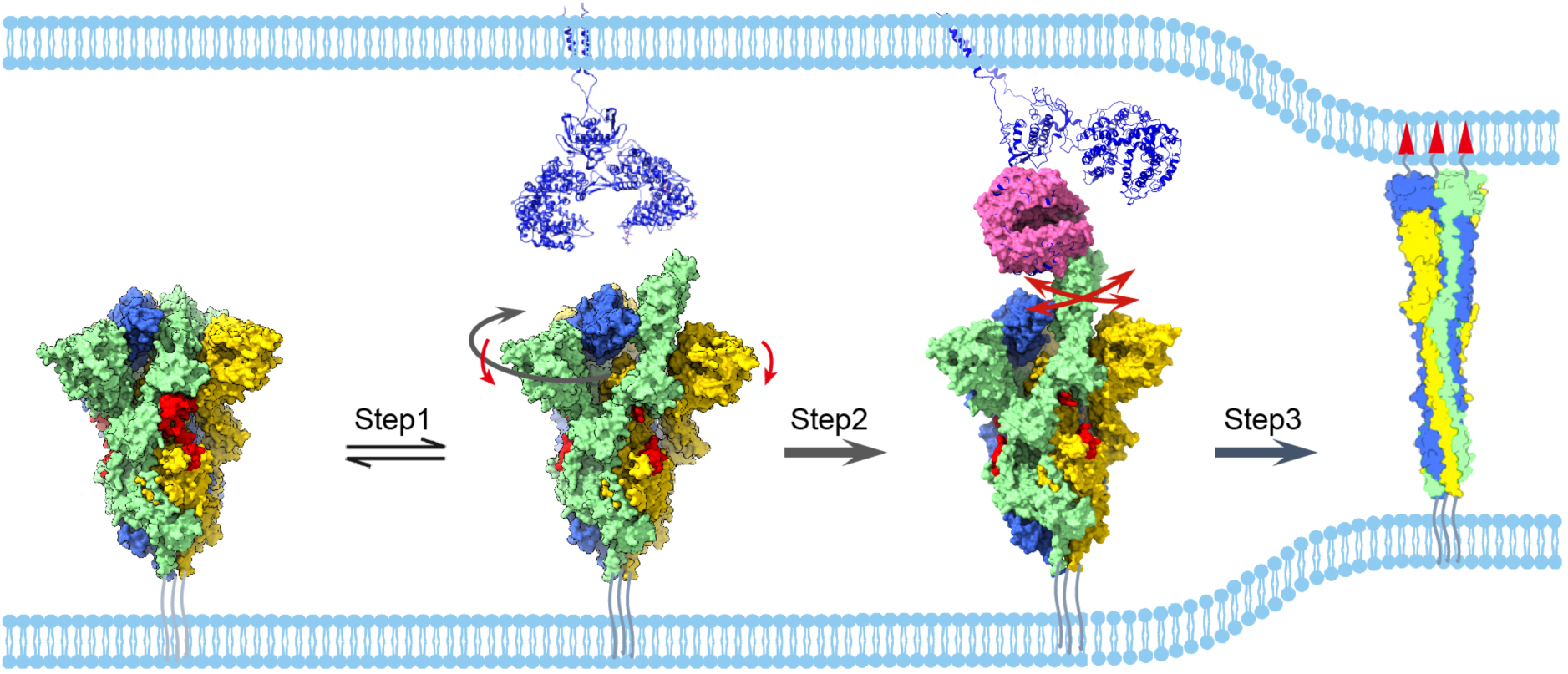
The proposed mechanism of ACE2 induced conformational transitions of SARS-CoV-2 S trimer. Conformational transitions from the ground prefusion closed state (with packed FP, in red) to the transiently open state (Step 1) with an untwisting motion of the S trimer (highlighted in dark grey arrow) associated with a downwards movement of S1 (red arrow), from the open state to the dynamic ACE2 engaged state (Step 2), and from the ACE2 engaged state all the way to the refolded postfusion state (Step 3). The continuous swing motions of the associated ACE2-RBD within the S trimer are indicated by red arrows. The SARS-CoV-2 S trimer associated with ACE2 dimer (in the third panel) was generated by aligning the ACE2 of our SARS-CoV-2 S-ACE2 complex structure with the available full length ACE2 dimer structure (PDB: 6M1D). The postfusion state was illustrated as a carton.

The dominantly populated conformation (94%) for the unliganded SARS-CoV-2 S trimer is in the tightly closed state (more compact than that of SARS-CoV S trimer) with all the RBD domains buried, resulting in conformational masking preventing antibody binding and neutralization at sites of receptor binding. This SARS-CoV-2 conformational masking mechanism of neutralization escape suggested here could affect all antibodies that bind to the receptor binding site, similar to that described for HIV-1 Env (Kwong et al., 2002; Munro et al., 2014). While for MERS-CoV or SARS-CoV S trimer, the closed state is less populated (5.4% and 27.6%, respectively, indicating the conformational masking mechanism may be less effective for the two viruses (Gui et al., 2017; Pallesen et al., 2017). Interestingly, our findings also suggest that unliganded S trimer proteins of SARS-CoV-2 are inherently competent to transiently display conformation with one RBD up ready for receptor ACE2 binding; ACE2 facilitates the capture of pre-existing S trimer open conformation that are spontaneously sampled in the unliganded spike, rather than triggering a trimer opening event. Therefore, the spontaneously sampled S trimer conformations may serve a functional role in infectivity.

Intriguingly, our data also suggest that the SARS-CoV-2 S trimer is very sensitive to ACE2. For instance, the presence of ACE2 triggers an extremely thorough conformational landscape transitions from dominantly closed state (94%) to all open configurations (including 26.2% free open state and 73.8% ACE2 bound open state). While in the counterpart of SARS-CoV, ACE2 induces conformational landscape transitions from 27.6% closed state and 72.4% open state to 24% closed state and 76% open state (including 26.7% free open state and 49.2% ACE2 bound open state) (Gui et al., 2017; Song et al., 2018). This demonstrates that the SARS-CoV-2 S trimer is much more sensitive to the ACE2 receptor than SARS-CoV S in terms of receptor-triggered transformation from the closed prefusion state to the fusion-prone open state, which might have contributed to the observed superior infectivity of SARS-CoV-2 as compared to that of SARS-CoV.

Noteworthy, the mutation SARS-CoV-2 spike D614G has gained urgent concern; the mutated genotype G614 began spreading in early February, and it was detected to reach at a frequency of ∼72% in early June according to GISAID public repository (Daniloski et al., 2020; Korber et al., 2020). Moreover, it has been reported that the D614G mutation promotes the infectivity of SARS-CoV-2 and enhances viral transmissibility in multiple human cell types (Daniloski et al., 2020; Hu et al., 2020; Zhang et al., 2020). However, the structural basis of D614G enhanced infectivity has not been fully understood yet. Here our S-closed structure in the ground prefusion state showed that D614 heavily involves in the interaction with FP through its side chain atoms (Fig. 1I, Table S2). This interaction could contribute greatly to the linkage between neighboring protomers as well as between S1 and S2 subunits. However, the mutation of D614 to G without side chain could eliminate most of the hydrogen bonds/salt bridges the D614 originally forms with FP, hence greatly reduce its interaction with FP potentially leading to a coordinated unpacking/activation of FPs. Therefore, D614G mutation could (1) reduce the constrains between neighboring protomers as well as S1/S2 interactions within S trimer, and (2) lower the energy barrier for the conformational transformation from closed prefusion state to fusion-prone open state, leading to even more sensitive SARS-CoV-2 S trimer to ACE2 binding. Collectively, these factors may contribute to the enhanced infectivity and viral transmissibility of the G614 strain.

In summary, our data revealed the unliganded SARS-CoV-2 S trimer to be intrinsically transforming between two distinct pre-fusion conformations, whose relative occupancies could be dramatically remodeled by receptor ACE2. These findings support a dynamics-based mechanism of immune evasion and ligand recognition (Munro et al., 2014). Thus, our study delineates the properties of the SARS-CoV-2 spike glycoproteins that simultaneously allow the retention of function and the evasion of the humoral immune response. We also delineated that the substantial conformational dynamics of S1 subunits induced by ACE2 binding could trigger the transition of the spike protein towards postfusion state prepared for viral entry and infection. Collectively, our findings suggest that stabilization of the tightly closed ground prefusion state of S trimer with inactivated FPs might be a general and effective means of inhibiting SARS-CoV-2 entry, and an understanding of the properties of the SARS-CoV-2 S trimer that permit neutralization resistance will guide attempts to create vaccines as well as therapeutics that target receptor binding.

## Acknowledgements

We are grateful to the staffs of the NCPSS Electron Microscopy facility, Database and Computing facility, and Protein Expression and Purification facility for instrument support and technical assistance. Y.C. was supported by grants from the CAS Pilot Strategic Science and Technology Projects B (XDB37040103), the National Basic Research Program of China (2017YFA0503503), the NSFC (31670754 and 31872714), the CAS Major Science and Technology Infrastructure Open Research Projects, and the CAS-Shanghai Science Research Center (CAS-SSRC-YH-2015-01, DSS-WXJZ-2018-0002). Z.H. was supported by grants from the Chinese Academy of Sciences (XDB29040300) and from the Ministry of Science and Technology of China (2020YFC0845900).

## Author contribution

Y.C. and Y.W. designed the experiments; Y.W., C.Z, Y.W., Y.Y. purified the proteins; X.H., Q.H., S.W., and C.L. performed NS-EM; C.L., X.H., and W.H. optimized the cryo-EM sample-preparation condition; C.L., W.H., Y.W. and W.Z. collected the cryo-EM data with the involvement of F.W. and L.K.; C.X. performed the cryo-EM reconstructions and the model building; C.X., C.L., X.H., Q.H., S.W., Q.Z., W.H., K.C. and Q.Z. performed particle picking; C.Z. and Y.W. performed the biochemical analyses; C.X., Y.C., Y.W., C.Z. and Z.H. analyzed the data, Y.C., Y.W., C.X. and Z.H. wrote the manuscript with inputs from all the others.

## Data deposition

Cryo-EM maps have been deposited in the Electron Microscopy Data Bank, https://www.ebi.ac.uk/pdbe/emdb/ (accession nos. ***), and the associated models have been deposited in the Protein Data Bank, www.rcsb.org (accession nos. **, **, and **).

## Conflict of interest statement

The authors declare that they have no conflict of interest.

## Methods

### Expression and purification of SARS-CoV-2 S trimer and human ACE2

To express SARS-CoV-2 S glycoprotein ectodomain, the mammalian codon-optimized gene coding SARS-CoV-2 (Wuhan-Hu-1 strain, GenBank ID: MN908947.3) S glycoprotein ectodomain (residues M1-Q1208) with proline substitutions at K986 and V987, a “GSAS” substitution at the furin cleavage site (R682–R685) was cloned into vector pcDNA 3.1+. A C-terminal T4 fibritin trimerization motif, a TEV protease cleavage site, a FLAG tag and a His tag were cloned downstream of the SARS-CoV-2 S glycoprotein ectodomain (Fig. S1A). A gene encoding human ACE2 PD domain (Q18-D615) with an N-terminal IL10 signal peptide and a C-terminal His tag was cloned into vector pcDNA 3.4. The expression vectors were transiently transfected into HEK293F cells using polyethylenimine. Three days after transfection, the supernatants were harvested. To purify the His-tagged S and ACE2 proteins, the clarified supernatants were added with 20 mM Tris-HCl pH 7.5, 200 mM NaCl, 20 mM imidazole, 4 mM MgCl_2_, and incubated with Ni-NTA resin at 4°C for 1 hour. The Ni-NTA resin was recovered and washed with 20 mM Tris-HCl pH 7.5, 200 mM NaCl, 20 mM imidazole. The protein was eluted by 20 mM Tris-HCl pH 7.5, 200 mM NaCl, 250 mM imidazole.

### Bio-layer interferometry (BLI) assay

Before BLI experiments, SARS-CoV-2 S trimer protein was biotinylated using the EZ-Link(tm) Sulfo-NHS-LC-LC-Biotin kit (Thermo Fisher) and then purified using Zeba(tm) spin desalting column (Thermo Fisher), according to manufacturer’s protocols. To determine binding affinity of ACE2, BLI assay was carried out using an Octet Red 96 instrument (Pall FortéBio, USA). Briefly, biotinylated SARS-CoV-2 S trimer protein was loaded onto streptavidin (SA) biosensors (Pall FortéBio). S-trimer-bound biosensors were dipped into wells containing varying concentrations of ACE2 protein and the interactions were monitored over a 500-sec association period. Finally, the sensors were switched to dissociation buffer (0.01 M PBS supplemented with 0.02% Tween 20 and 0.1% bovine serum albumin) for a 500-sec dissociation phase. Data was analyzed using Octet data analysis software version 11.0 (Pall FortéBio).

### SARS-CoV-2 S-ACE2 complex formation

The purified SARS-CoV-2 S glycoprotein ectodomain and human ACE2 PD domain were mixed at a molar ratio of 1:3 and were incubated on ice for 2 hours. The mixture was purified by filtration chromatography using a Superose 6 increase 10/300 GL column (GE Healthcare) pre-equilibrated with 20 mM Tris-HCl pH 7.5, 200 mM NaCl, 4% glycerol. For cross linking complex, the buffer of purified SARS-CoV-2 S glycoprotein ectodomain and human ACE2 PD domain were exchanged to 20 mM Hepes pH 7.5, 200 mM NaCl; then SARS-CoV-2 S and human ACE2 were mixed at a molar ratio of 1:3. After incubation on ice for 2 hours, the complex was cross linked by 0.1% glutaraldehyde, which is commonly used in cryo-EM studies of fragile macromolecular complexes (Kastner et al., 2008; Patel et al., 2018). The glutaraldehyde was neutralized by adding 20 mM Tris-HCl pH 7.5 after incubated on ice for 1 hour. The mixture was run over a Superose 6 increase 10/300 GL column (GE Healthcare) in 20 mM Tris-HCl pH 7.5, 200 mM NaCl, 4% glycerol. The complex peak fractions were concentrated and assessed by SDS-PAGE and negative-staining electron microscopy.

### Negative-stain sample preparation, data collection and initial model building

For the NS sample, a volume of 5 µL of SARS-CoV-2 S-ACE2 sample was placed on a plasma cleaned copper grid for one minute. Excess sample on the grid was blotted off using filter paper, and a volume of 5 µl of 0.75% UF (Sigma-Aldrich) was added to wash the grid. After blotting, another volume of 5 µL of 0.75% UF was placed on the grid again for one minute to stain. Grids were visualized under a Tecnai G2 Spirit 120 kV transmission electron microscope (Thermo Fisher Scientific), and micrographs were taken using an Eagle camera with a nominal magnification of 67,000×, yielding a pixel size of 1.74 Å. 41,827 particles were autopicked in EMAN2 (Bell et al., 2016). After 2D classification, we selected good averages with 13,047 particles for initial model building, which were performed in RELION 3.0 (Zivanov et al., 2018).

### Cryo-EM sample preparation for SARS-CoV-2 S trimer and S-ACE2 complex

To prepare the cryo-EM sample of SARS-CoV-2 S trimer, a 2.2-µL aliquot of this sample was applied to a plasma cleaned holey carbon grid (R2/1, 200 mesh; Quantifoil) or Graphene Oxide-Lacey Carbon grid (300 mesh, EMR). The grid was blotted with Vitrobot Mark IV (Thermo Fisher Scientific) and then plunged into liquid ethane cooled by liquid nitrogen. To prepare the cryo-EM sample of S-ACE2 complex with or without cross linking, we used Graphene Oxide-Lacey Carbon grid (300 mesh, EMR), and adopted the same vitrification procedure as for the S trimer.

### Cryo-EM data collection

Cryo-EM movies of the samples were collected on a Titan Krios electron microscope (Thermo Fisher Scientific) operated at an accelerating voltage of 300 kV with a nominal magnification of 22,500x (Table S1). The movies were recorded on a K2 Summit direct electron detector (Gatan) operated in the super-resolution mode (yielding a pixel size of 1.02 Å after 2 times binning), under low-dose condition in an automatic manner using SerialEM (Mastronarde, 2005). Each frame was exposed for 0.15 s and the total accumulation time was 6.45s, leading to a total accumulated dose of 50 e^-^/Å^2^ on the specimen. To solve the problem of preferred orientation associated with SARS-CoV-2 S trimer, we additionally collected tilt datasets with the stage tilt at 30° or 40°, while the other conditions remained the same.

### Cryo-EM 3D reconstruction

Single particle analysis was mainly executed in RELION 3.1 (Fernandez-Leiro and Scheres, 2017). All images were aligned and summed using MotionCor2 software (Zheng et al., 2017). After CTF parameter determination using CTFFIND4 (Rohou and Grigorieff, 2015), particle auto-picking, manual particle checking, and reference-free 2D classification, particles with S trimer features were maintained for further processing.

For receptor-free S trimer sample, 226,082 particles were picked from non-tilt micrographs, and 118,420 remained after 2D classification (Fig. S3). These particles went through 3D auto-refine using available SARS-CoV-2 S trimer cryo-EM map (EMDB: 21452) lowpass filtered to 40 Å resolution as initial model (Walls et al., 2020). These particles were refined into a closed state map of S trimer with imposed C3 symmetry. We then re-extracted the particles using the refinement coordinates to re-center it. After CTF refinement and polishing, these particles were refined with C3 symmetry again. Noteworthy, the Euler angle distribution of the map suggested the dataset is lacking tilted top views (Fig. S2C left panel). Indeed, when refine the dataset without imposing 3-fold symmetry, the top view of the map appeared distorted indicating a preferred orientation problem associated with the sample. To overcome the preferred orientation problem, we additionally collected tilt data, and boxed out 198,737 particles from 40° tilt micrographs and 16,010 particles from 30° tilt micrographs. After 2D classification, 184,661 particles remained. We then used goCTF software to determine the defocus for each of the tilt particle, and these particles were re-extracted with corrected defocus (Su, 2019). After combining the tilt with non-tilt particles, we refined the dataset without imposing symmetry, then performed two rounds of 3D and 2D classifications to further cleanup the dataset, and obtained a dataset of 151,505 particles, of which 62,368 particles were from the tilt data. We then carried out heterogeneous refinement in CryoSparc (Punjani et al., 2017), and obtained a closed state map from 142,345 particles and an open state reconstruction with 9,160 particles (Fig S3). After CTF refinement and Bayesian polishing, the closed state map was refined to 2.7 Å resolution with C3 symmetry, while the open state map was at 12.8 Å resolution and hardly to improve the resolution, indicating an intrinsic dynamic nature of the open state. The overall resolution was determined based on the gold-standard criterion using an FSC of 0.143 (Scheres and Chen, 2012).

For the SARS-CoV-2 S-ACE2 cross-linked dataset, 298,127 particles were picked from original micrographs, and 138,632 particles remained after 2D classification (Fig. S5). These particles were refined with an initial model built from our negative staining data. We then re-extracted the particles to re-center them. These particles went through a 3D-2D classification step resulting in a further cleaned up dataset of 77,440 particles. We refined these particles into a map of ACE2 bound S trimer complex. We then used this map as initial model to refine the originally picked 298,127 particles for one round to re-extract and re-center the particles. After 2D classification, 207,742 particles remained. After 2 rounds of 3D-2D cleaning step, 136,412 particles were left for further structure determination. After heterogeneous refinement in CryoSparc, class 1 resembled an ACE2-free open state of S trimer, and classes 2-5 adopted S-ACE2 engaged conformation. For class 1, after further 2D classification, we refined the 24,502 cleaned up particles into a S-open map at 6.0 Å resolution using non-uniform refinement in CryoSparc. Among the other four classes with bound ACE2, we sorted out good particles for classes 2-4 by 2D classification and combined them with class 5 exhibiting good structural details, resulting in a dataset of 68,987 particles. After refinement, Bayesian Polishing, and CTF refinement, we reconstructed a 3.8 Å resolution SARS-CoV-2 S-ACE2 map. The S trimer portion without the up RBD was rather stable, could be locally refined to 3.3 Å using local refinement in CryoSparc with non-uniform refinement option chosen. The ACE2 associated with the up RBD was subtracted and refined in Relion to obtain a more 8.4 Å map with better connectivity. Multi-body refinement in Relion 3.1 was applied to analyze the motion of the complex.

For SARS-CoV-2 S-ACE2 w/o crosslinking dataset, we followed similar classification and cleaning up strategy and obtained 81,820 particles. Through heterogeneous refinement and 2D classification in CryoSparc, we reconstructed a 5.3 Å resolution SARS-CoV-2 S-ACE2 map from 32,866 particles using non-uniform refinement, and an unliganded open state map of 11.2 Å resolution from 15,149 particles, with the population of 68.4% and 31.6%, respectively. Multi-body refinement was also applied to analysis the mobility of the complex.

### Pseudo atomic model building

To build the pseudo atomic model for our SARS-CoV-2 S-closed structure, we used the available atomic model of SARS-CoV-2 S (PDB: 6VXX) as initial model (Walls et al., 2020). We first refined the model against our map using phenix.real_space_refine module in Phenix (Adams et al., 2010). For the missing loop regions in S1 subunit, we either built the homology model based on SARS-CoV S structure (PDB: 6CRW) (Kirchdoerfer et al., 2018) through SWISS-MODEL webserver (Waterhouse et al., 2018), or built the loop manually according to the density in COOT (Emsley and Cowtan, 2004). For the FP region, we first built the homology model by Modeller tool within Chimera by using MERS-CoV S structure (PDB: 6NB3) as template (Pettersen et al., 2004; Sali, 1995; Walls et al., 2019), then used Rosetta to refine this region against the density map (DiMaio et al., 2015). Eventually, we used phenix.real_space_refine again for the protomer and S-trimer model refinement against the map.

For the SARS-CoV-2 S-ACE2 structure, we used the SARS-CoV-2 RBD-ACE2 crystal structure (PDB: 6M0J) as initial model for the ACE2 and the associated up RBD portion, and our S-closed model as initial model for the remaining portion. These models were firstly refined against the corresponding focused map using Rosetta and Phenix (DiMaio et al., 2015), then combined together in COOT. We then refined the combined model against our 3.8 Å resolution SARS-CoV-2 S-ACE2 map using Rosetta and Phenix. For the S-open structure, we used the model of SARS-CoV-2 S-ACE2 as initial model with ACE2 removed, and refined against the map using Rosetta.

We used Phenix.molprobility to evaluate the models, and calculated B-factors by atom displacement refinement function in phenix.real_space_refine. We used UCSF Chimera and ChimeraX for figure generation (Goddard et al., 2018; Pettersen et al., 2004), and also for rotation, translation, RMSD, and vdw contact measurement. Interaction surface analysis was conducted by PISA server (Krissinel and Henrick, 2007).

### Identification of key amino acids involved in ACE2 recognition with RBD mutants by ELISA

To uncover the amino acids important for ACE2 receptor recognition, ACE2 ecotodomain (residues Q18 to S740) gene, with an N-terminal IL10 signal peptide, tagged with human IgG1Fc and His tag at the C-terminus, were cloned into the pcDNA 3.4 vector. Codon-optimized RBD (residues V320 to G550) gene fragment, with an N-terminal IL10 signal peptide, tagged with His tag at the C-terminus, were cloned into the pcDNA 3.4 vector. Three SARS-CoV-2 RBD mutants were constructed. For mutant RBD (Core), amino acids R319 to N437 of core region in the SARS-CoV-2 RBD were substituted by the corresponding region of SARS-CoV strain Tor2 (GenBank ID: AAP41037.1). For mutants RBD (RBM-R2) and RBD (RBM-R3), residues L452 to K462, and residues T470 to T478 of RBM region in the SARS-CoV-2 RBD were mutated into the corresponding regions of SARS-CoV strain Tor2, respectively. For single point mutations of RBD (Q498A), RBD (V503A), and RBD (Y505A), RBD residues Q498, V503, and Y505 were substituted by Ala, respectively. All mutant plasmids were constructed using the MutExpress™ II Fast Mutagenesis Kit V2 (Vazyme, China) according to the manufacturer’s instruction. The proteins were generated using HEK 293F expression system and purified as described above.

Anti-RBD polyclonal antibody and monoclonal antibody (MAb) 1A10 were prepared by immunizing BALB/c mice with recombinant SARS-CoV-2 RBD fused with a C-terminal mouse IgGFc tag (Sino Biological Inc, Beijing, China) using previously described protocols (Qu et al., 2020).

The purified RBD mutants were tested by ELISA for reactivity with the receptor ACE2. Briefly, ELISA plates were coated with 100 ng/well of the purified RBD mutants in PBS at 37°C for 2 hours and then blocked with 5% milk in PBS-Tween20 (PBST). Next, the plates were incubated with 50 ng/well of ACE2-hFc fusion protein, 50 µL/well of culture supernatant of hybridoma 1A10, or 50 µL/well of mouse anti-RBD sera (diluted at 1/1000) at 37°C for 2 h. After washing, the corresponding secondary antibodies, horseradish peroxidase (HRP)-conjugated anti-human IgG1 (Abcam, USA) or HRP-conjugated anti-mouse IgG (Sigma, USA), were added and incubated at 37°C for 1 h. After washing color development, absorbance at 450 nm was determined.

## Supplementary Information

**Figure S1.**
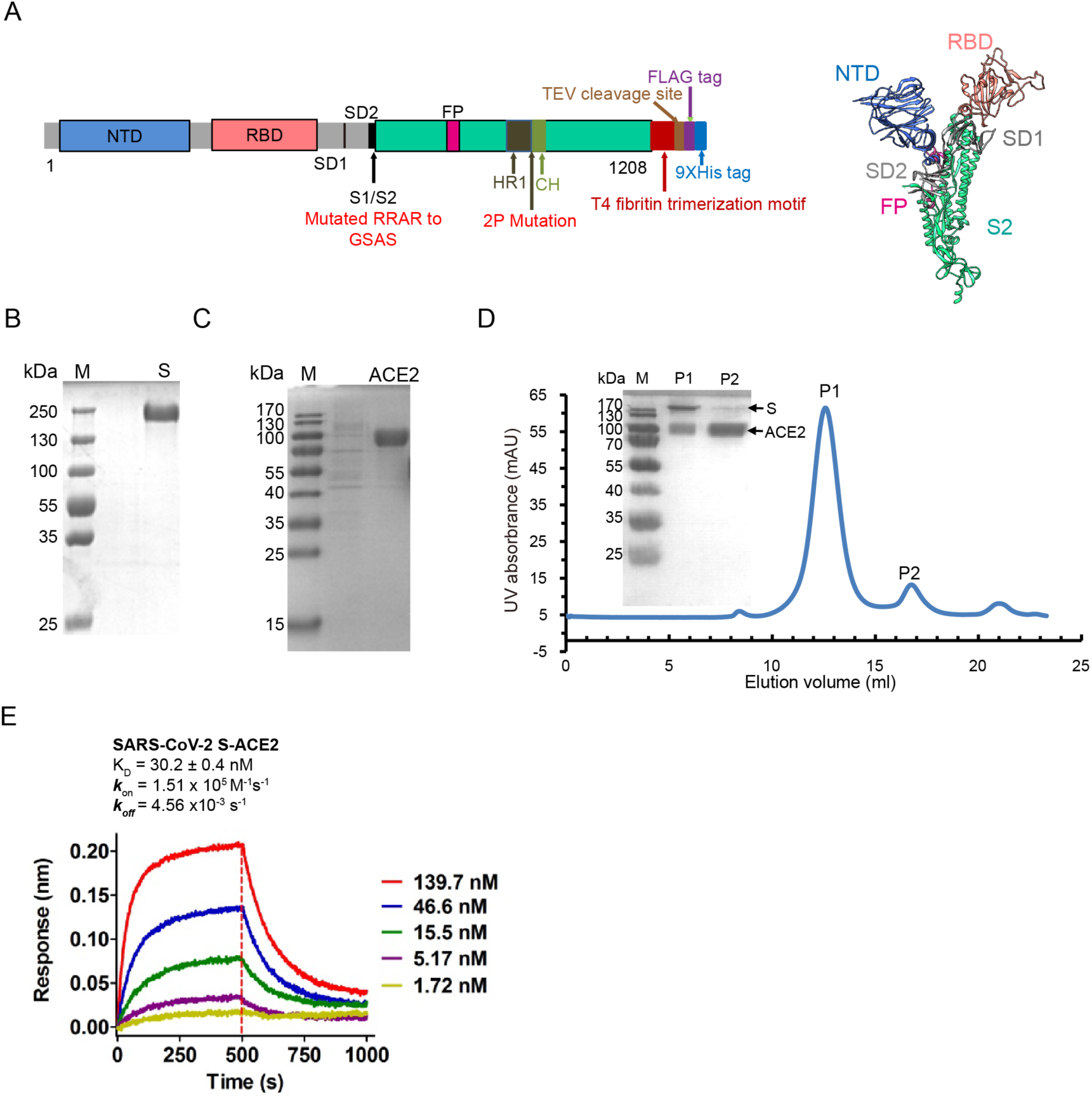
Purification of SARS-CoV-2 S ectodomain, human ACE2 PD domain, and SARS-CoV-2 S-ACE2 complex. (A) Schematic diagram of SARS-CoV-2 S organization in this study. S1/S2 protease cleavage site (S1/S2), N-terminal domain (NTD), receptor-binding domain (RBD), fusion peptide (FP), heptad repeat 1 (HR1), and central helix (CH) are labeled. (B-C) SDS-PAGE analysis of the purified S protein (B) and ACE2 (C). (D) Size-exclusion chromatogram and SDS-PAGE analysis of the formed SARS-CoV-2 S-ACE2 complex.

**Figure S2.**
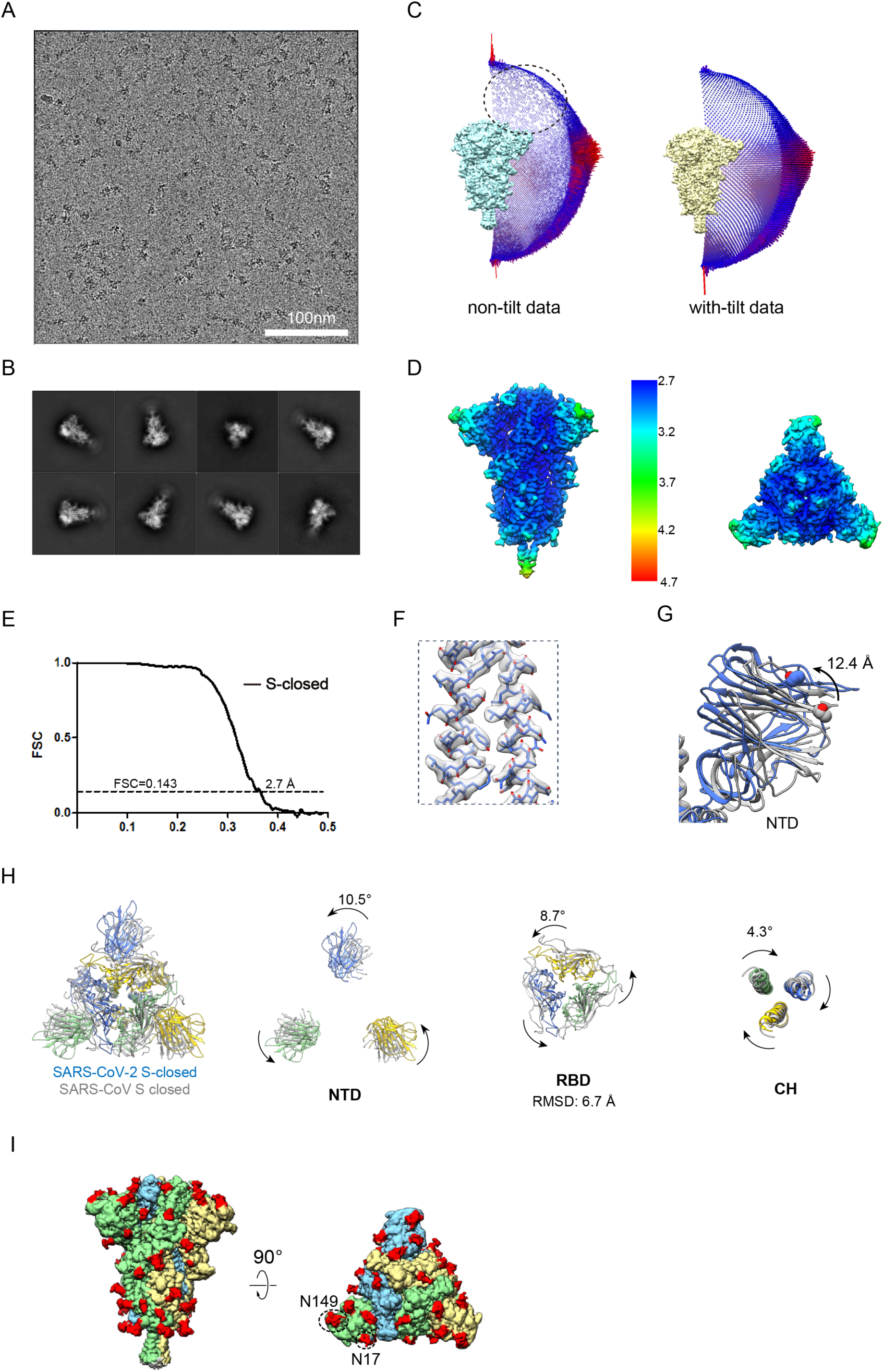
Cryo-EM analysis of the SARS-CoV-2 S trimer in the tightly closed state. (A) Representative cryo-EM image of the SARS-CoV-2 S trimer. (B) Reference-free 2D class averages of the S trimer. (C) Euler angular distribution of 3D reconstructions before and after adding tilt data, with the dotted circle indicating the sparsely distributed tilted top views in the non-tilt data. (D-E) Local resolution evaluation (D) and resolution assessment of our S-closed cryo-EM map by Fourier shell correlation (FSC) at 0.143 criterion (E). (F) Close up view of the model-map fitting in S2 subunit. (G) Compared with the recent structure of SARS-CoV-2 S in closed state (gray, 6VXX), our S-closed structure (blue) showed a slight inward tilt leading the peripheral edge of NTD exhibiting a 12.4 Å inward movement (for the Cα of T124). (H) Top view of the overlaid structures between our SARS-CoV-2 S-closed structure and the SARS-CoV S-closed structure (PDB: 5XLR) and zoom in views of the overlaid structures in NTD, RBD, CH domains. (I) N-linked glycans resolved in our S-closed cryo-EM map, with the densities corresponding to glycans colored in red.

**Figure S3.**
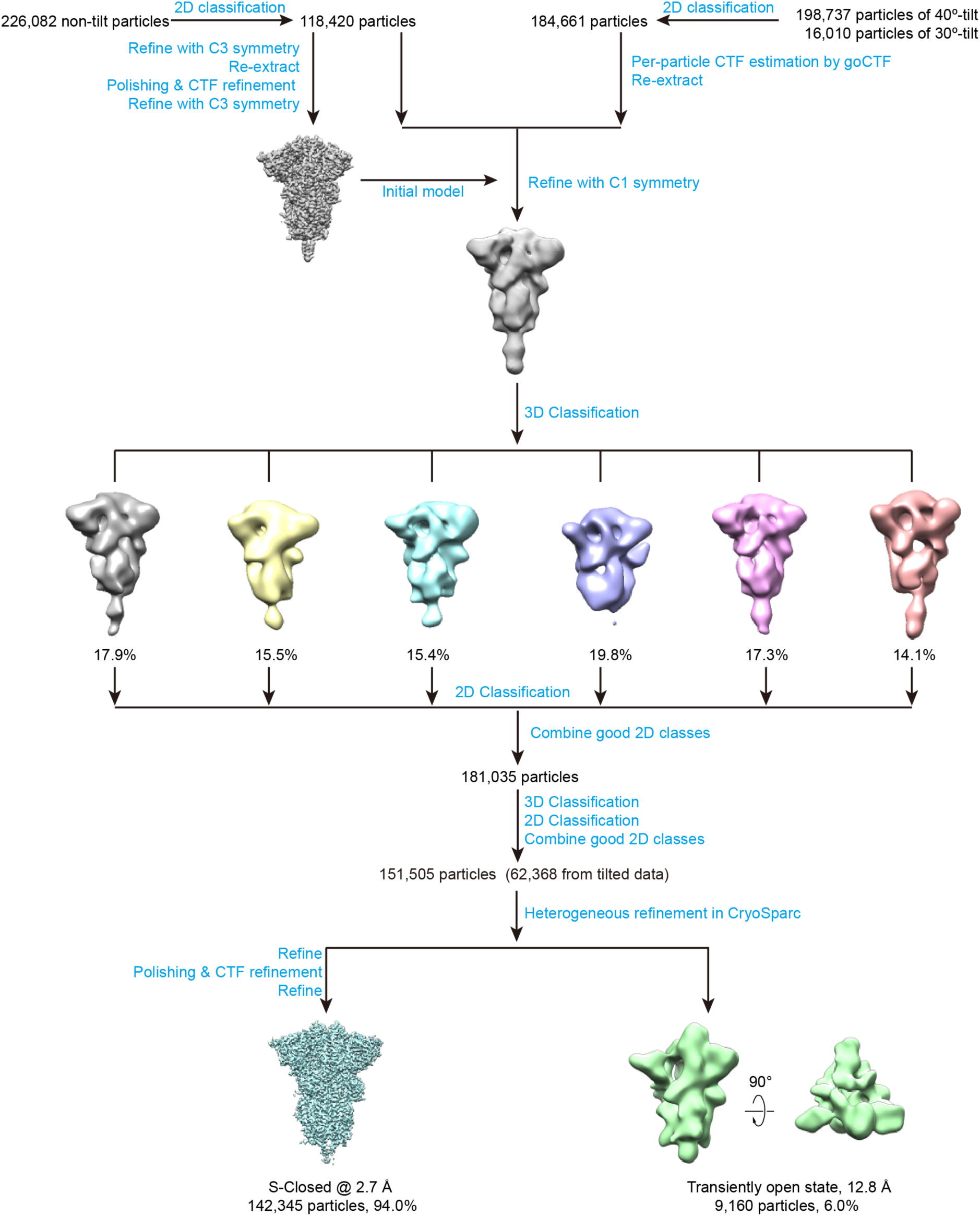
Cryo-EM data processing procedure for SARS-CoV-2 S trimer.

**Figure S4.**
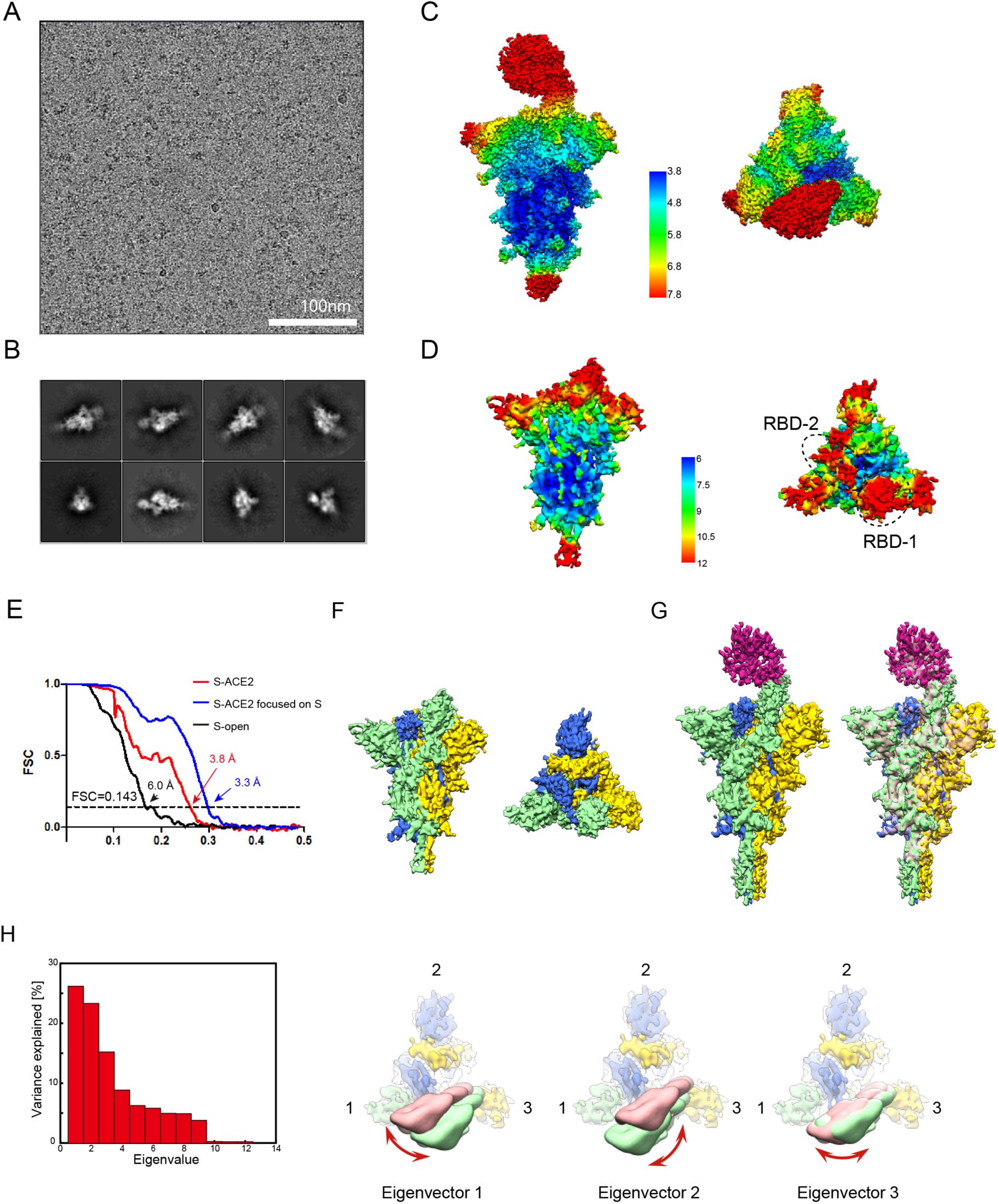
Cryo-EM analysis on the SARS-CoV-2 S-ACE2 complex. (A) Representative cryo-EM image of the SARS-CoV-2 S trimer in the presence of ACE2. (B) Reference-free 2D class averages of the sample. (C-D) Local resolution evaluation of the S-ACE2 map (C) and S-open map (D). (E) Resolution assessment of the cryo-EM reconstructions by Fourier shell correlation (FSC) at 0.143 criterion. (F) Unliganded S-open map obtained from this dataset. (G) Cryo-EM map of S-ACE2 complex without cross linker (left, colored), and its overlay with S-ACE2 map with cross linker (pink, low pass filtered to similar resolution, right panel), suggesting they are in similar conformation. (H) ACE2 binding induced motions of S-ACE2 without cross linker. Left, contributions of all eigenvectors to motions of S-ACE2; right three panels, top view of the map showing the three swing motions along the first 3 eigenvectors.

**Figure S5.**
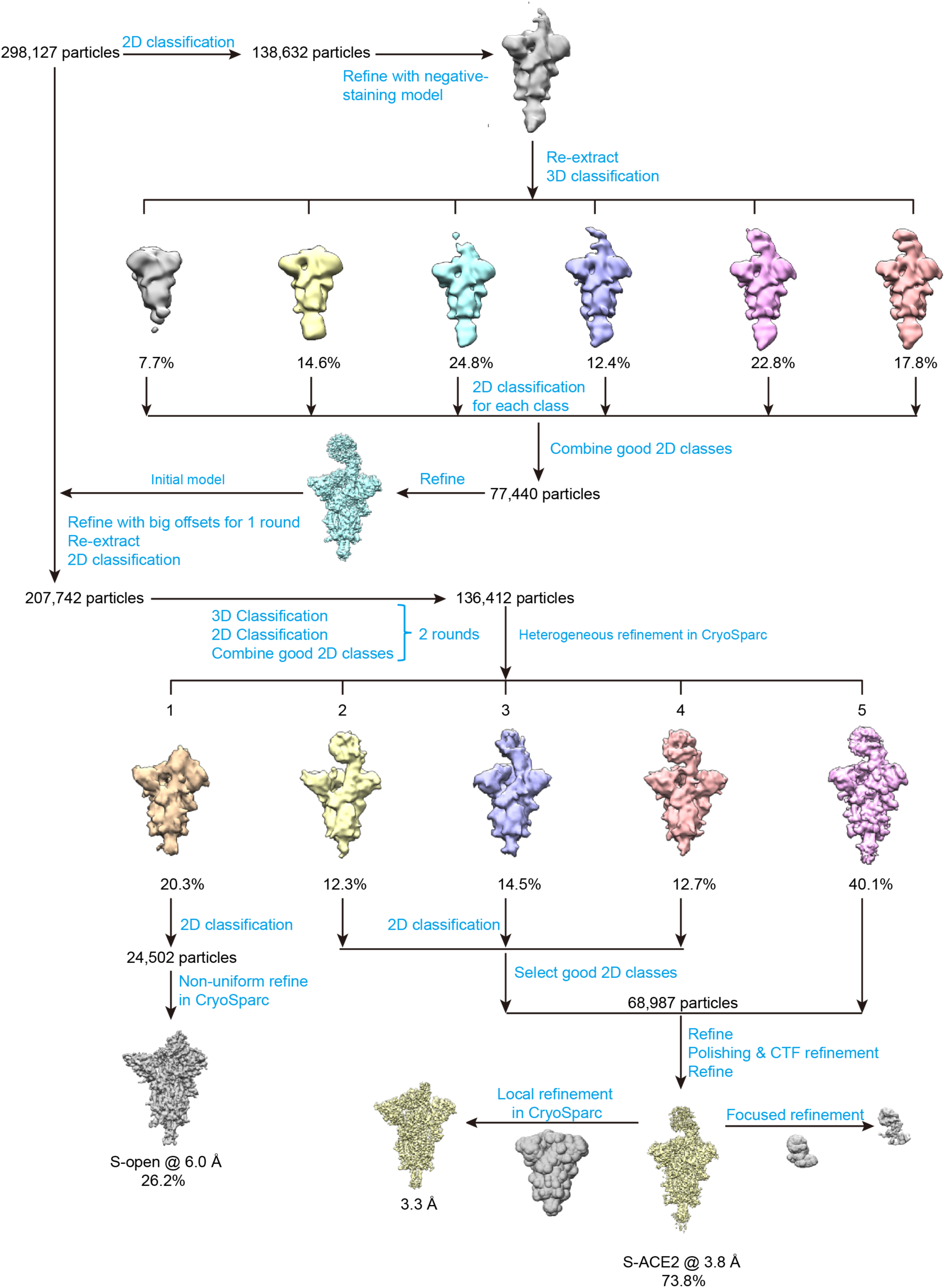
Cryo-EM data processing procedure for SARS-CoV-2 S trimer in the presence of ACE2.

**Figure S6.**
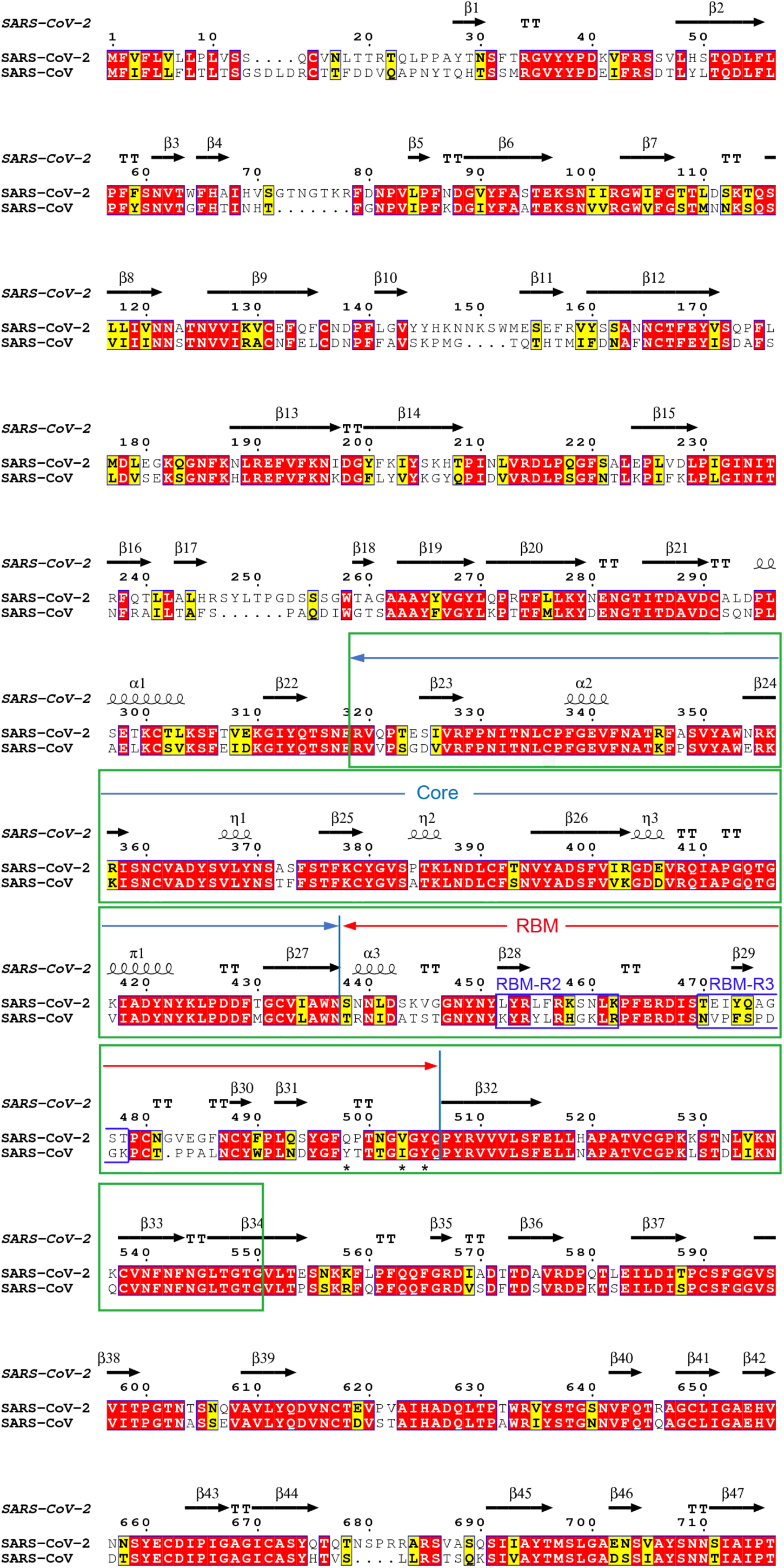
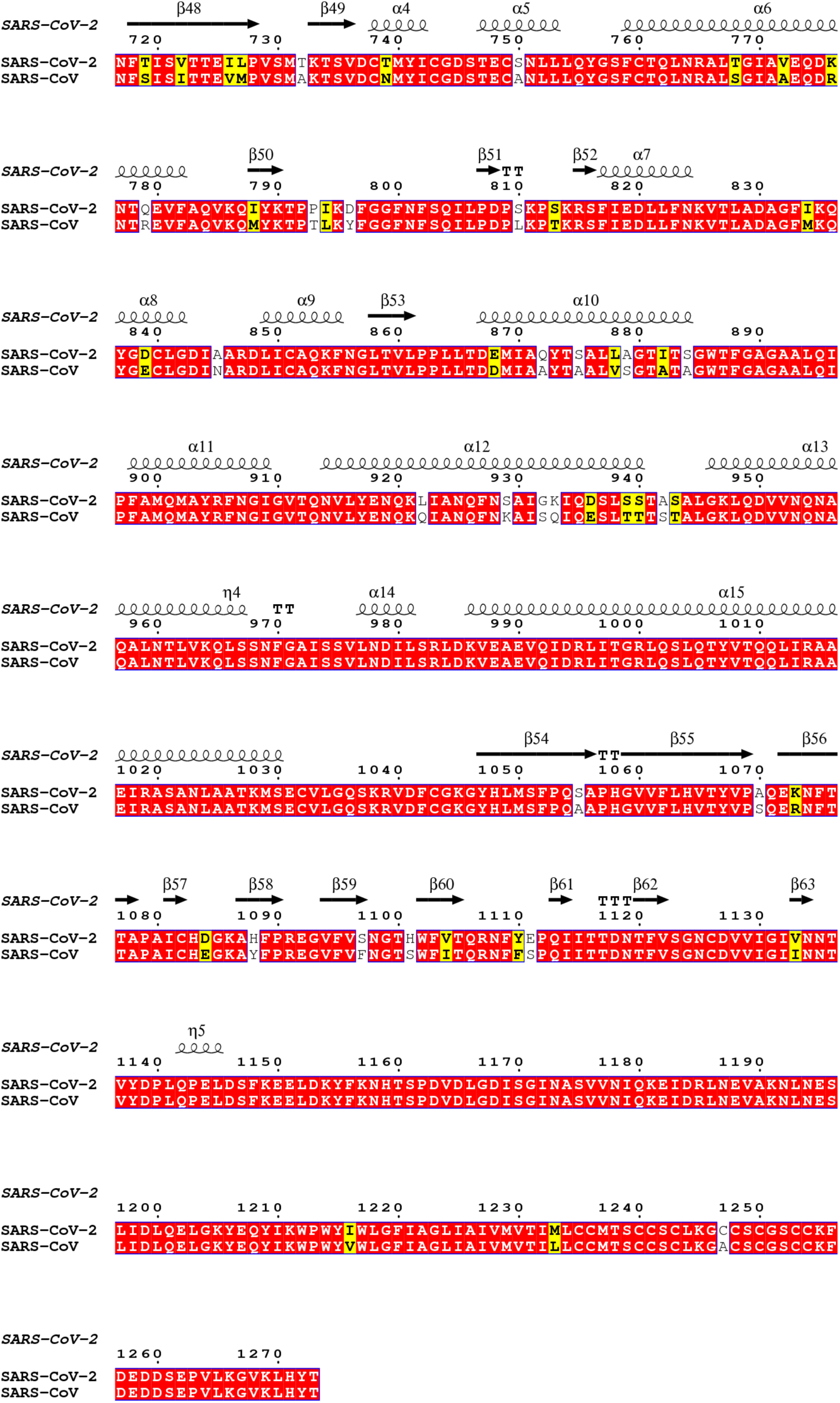
Amino acid sequence alignment of SARS-CoV-2 S to SARS-CoV S. The secondary structure elements were defined based on an ESPript (Robert and Gouet, 2014) algorithm and are labeled based on our SARS-CoV-2 S-closed structure. The RBD domain is labeled in green frames, and the subdomains of RBM are also labeled.

**Table S1.**
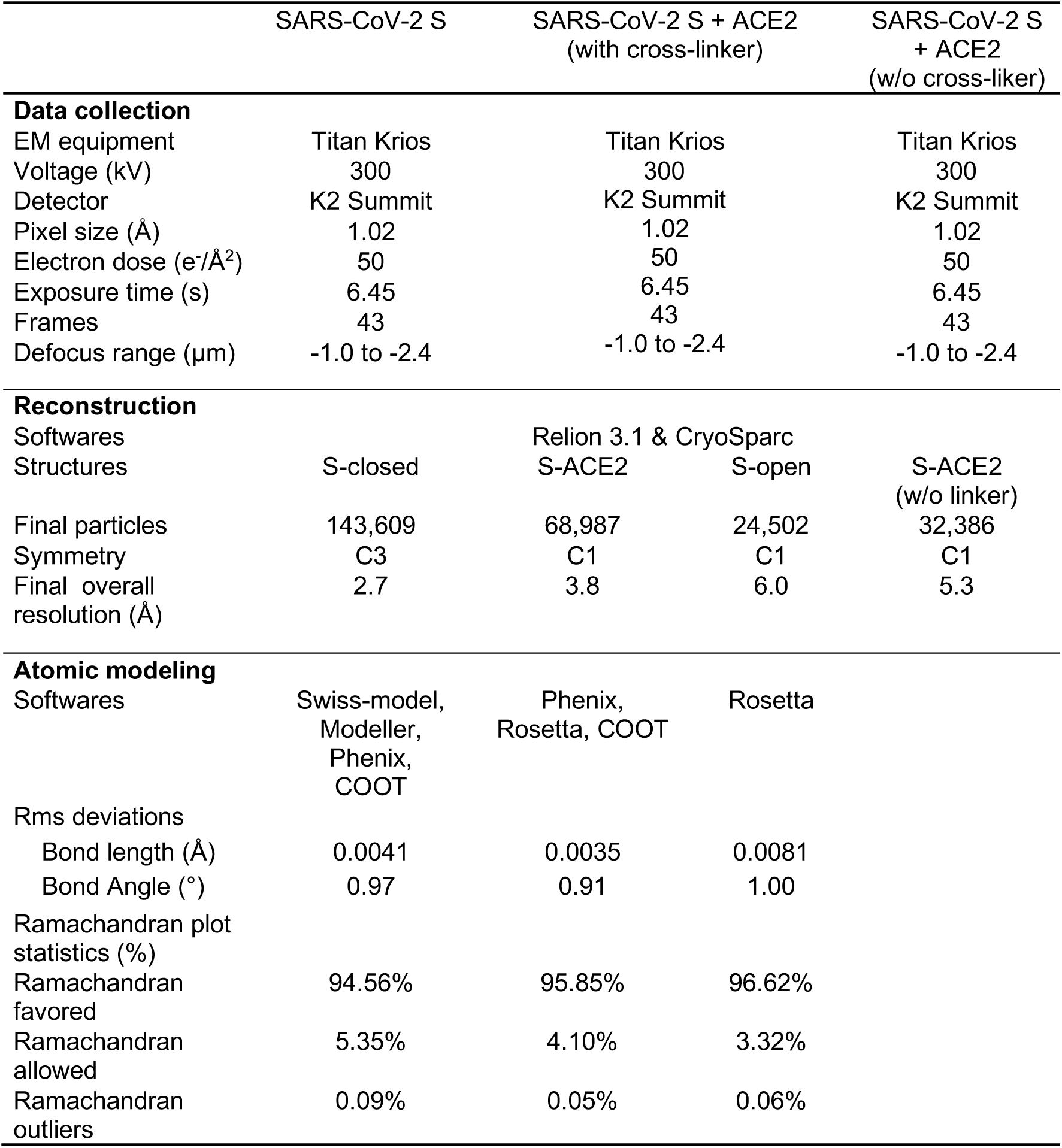
Cryo-EM data collection and refinement statistics.

**Table S2.** Fusion peptide D828-F855 region involved interactions

| D828-F855<br>region (FP) |  | Other motifs |  |  | Interaction | Distance(Å) |
| --- | --- | --- | --- | --- | --- | --- |
| Residue | Atom | Residue | Atom | Location |  |  |
| K835 | N | D614 | OD1 | SD2 (prot 1) | H-bond | 3.74 |
| Y837 | N | D614 | O | SD2 (prot 1) | H-bond | 3.89 |
| Y837 | OH | P589 | O | SD2 (prot 1) | H-bond | 3.56 |
| K854 | NZ | D614 | OD2 | SD2 (prot 1) | Salt bridge<br>/H-bond | 2.67 |
| K854 | NZ | D614 | OD1 | SD2 (prot 1) | Salt bridge | 3.18 |
| Q836 | OE1 | N616 | N | SD2 (prot 1) | H-bond | 2.72 |
| Q853 | NE2 | A956 | O | HR1 (prot 3) | H-bond | 3.84 |
| Q853 | NE2 | N960 | OD1 | HR1 (prot 3) | H-bond | 2.35 |

**Table S3.** SARS-CoV-2 S-ACE2 structure revealed RBD-ACE2 interactions

RBD-ACE2 H-bonds
| RBD from SARS-CoV-2 S |  | ACE2 |  | Interaction | Distance(Å) |
| --- | --- | --- | --- | --- | --- |
| Residue | Atom | Residue | Atom |  |  |
| G446 | O | Q42 | NE2 | H-bond | 3.49 |
| A475 | O | S19 | N | H-bond | 2.43 |
| N487 | OD1 | Y83 | OH | H-bond | 2.26 |
| Y489 | OH | Y83 | OH | H-bond | 3.72 |
| S494 | O | H84 | NE2 | H-bond | 2.42 |
| T500 | OG1 | Y41 | OH | H-bond | 2.45 |
| G502 | N | K353 | O | H-bond | 2.44 |
| Y505 | OH | A386 | O | H-bond | 2.36 |
| Y505 | OH | R393 | NH2 | H-bond | 2.43 |

Contacting residues at the SARS-CoV-2 RBD-ACE2 interface (distance cutoff of 4 Å)
| SARS-CoV-2 S RBD | ACE2 |
| --- | --- |
| Y453 | H34 |
| L455 | K31 |
| F456 | T27, D30, K31 |
| Y473 | T27 |
| A475 | S19, T27 |
| G476 | S19, Q24 |
| F486 | L79, M82, Y83 |
| N487 | Q24, Y83 |
| Y489 | T27, F28, K31 |
| F490 | K31 |
| Q493 | K31, H34 |
| S494 | H34 |
| Y495 | H34 |
| G496 | K353 |
| Q498 | Y41 |
| T500 | Y41, D355, R357 |
| N501 | K353 |
| G502 | K353, G354 |
| Y505 | K353, G354, A386, R393 |

## Notes

### Competing Interest Statement

The authors have declared no competing interest.

